# Dopamine D2Rs Coordinate Cue-Evoked Changes in Striatal Acetylcholine Levels

**DOI:** 10.1101/2021.12.08.471871

**Authors:** Kelly M. Martyniuk, Arturo Torres-Herraez, Marcelo Rubinstein, Marie A. Labouesse, Christoph Kellendonk

## Abstract

In the striatum, acetylcholine (ACh) neuron activity is modulated co-incident with dopamine (DA) release in response to unpredicted rewards and reward predicting cues and both neuromodulators are thought to regulate each other. While this co-regulation has been studied using stimulation studies, the existence of this mutual regulation *in vivo* during natural behavior is still largely unexplored. One long-standing controversy has been whether striatal DA is responsible for the induction of the cholinergic pause or whether D2R modulate a pause that is induced by other mechanisms. Here, we used genetically encoded sensors in combination with pharmacological and genetic inactivation of D2Rs from cholinergic interneurons (CINs) to simultaneously measure ACh and DA levels after CIN D2R inactivation. We found that CIN D2Rs are not necessary for the induction of cue induced dips in ACh levels but regulate dip lengths and rebound ACh levels. Importantly, D2R inactivation strongly decreased the temporal correlation between DA and Ach signals not only at cue presentation but also during the intertrial interval. This points to a general mechanism by which D2Rs coordinate both signals. At the behavioral level D2R antagonism increased the latency to lever press, which was not observed in CIN-selective D2R knock out mice. This latency correlated with the cue evoked dip length supporting a role of the ACh dip and it’s regulation by D2Rs in motivated behavior. Overall, our data indicate that striatal DA coordinate phasic ACh and DA signals via CIN D2Rs which is important for the regulation of motivated behavior.

## Introduction

Dopamine (DA) plays a key role in learning, serving as a teaching signal that reflects reward prediction error (Day et al., 2007; Mohebi et al., 2019; Nasser et al., 2017; Schultz et al., 1997; Steinberg et al., 2013). This teaching function is encoded in the phasic bursting of DA neurons, which induces a rapid but transient increase of extracellular DA. DA is initially released in response to an unpredicted reward, but with learning the response shifts away from the reward outcome towards reward predicting cues (Schultz, 2007; Schultz et al., 1997).

Like DA neurons, cholinergic interneurons (CINs) in rodents and their presumed counterparts, “tonically active neurons” (TANs), in primates modulate their activity in response to reward predicting cues and salient outcomes. CINs represent about 1-2% of the neurons in the striatum and regulate mental processes including reinforcement learning, action selection, associative learning, and cognitive flexibility (Aoki et al., 2015; Bradfield et al., 2013; Joshua et al., 2008; Matamales et al., 2016; Maurice et al., 2015; Morris et al., 2004; Okada et al., 2014). Pharmacogenetic inhibition of CINs in the NAc also increases the influence of appetitive cues on instrumental actions pointing to a role of striatal ACh in motivation (Collins et al., 2019). CINs are tonically active and show a multiphasic response to salient and conditioned stimuli that can include a short excitation followed by a prominent pause and rebound excitation (Aosaki et al., 1994a; Aosaki T, 1994b; Apicella, 2007; Apicella et al., 2009; Apicella et al., 2011). This multiphasic response in CIN firing coincides with phasic activation of midbrain DA neurons that terminate in the striatum (Joshua et al., 2008; Morris et al., 2004; Schultz, 2007; Schultz et al., 1997). Furthermore, there is increasing evidence that DA and ACh regulate each other within the striatum (Cachope & Cheer, 2014; Cachope et al., 2012; Chuhma et al., 2014; Cragg, 2006; Helseth AR, 2021; Kharkwal et al., 2016; Straub et al., 2014; Sulzer et al., 2016; Threlfell et al., 2012; Yan & Surmeier, 1991).

Here, we will focus on the DA-regulation of the multiphasic ACh response. One long-standing discussion in this regard has been whether the cholinergic pause is dependent on DA via DA D2R receptor (D2R) mediated inhibition of CINs. Early evidence that the CIN pause is DA dependent originate from studies in non-human primates (NHPs). *In vivo* electrophysiological recordings from TANs have revealed a pronounced pause in firing to a reward-predicting stimulus. This pause was entirely abolished by 1-methyl-4-phenyl-1,2,3,6-tetrahydropyridine (MPTP) lesions of DA neurons and local administration of a D2R antagonist (Aosaki et al., 1994a; Watanabe & Kimura, 1998). Consistent with this, more recent slice physiology studies in rodents have shown that pauses in CIN activity can be induced by local application of DA or DA terminal stimulation, which both are eliminated by pharmacological blockade of D2Rs (Augustin et al., 2018; Chuhma et al., 2014; Straub et al., 2014; Wieland et al., 2014). Additionally, optogenetic stimulation of NAc DA terminals results in a pause in CIN firing and this pause is prolonged when D2Rs are selectively overexpressed in CINs (Gallo et al., 2021). Lastly, pauses generated by DA or local stimulation of the striatum are eliminated in a selective CIN D2 knockout mouse (Augustin et al., 2018; Kharkwal et al., 2016). Taken together, the slice physiology experiments provide evidence that the CIN pause can be induced by DA activation in a CIN D2R dependent manner while the NHP studies show the necessity for DA and D2Rs for the generation of the pause.

However, more recent evidence suggests that the CIN pause is not induced by DA but by cortical, thalamic, or long-range GABAergic inputs (Brown et al., 2012; Cover et al., 2019; Ding et al., 2010; Doig et al., 2014; English et al., 2012; Matsumoto et al., 2000; Zhang et al., 2018). Consistent with this, stimulation of cortical and thalamic inputs to the striatum in slices or *in vivo* induces a triphasic cholinergic pause. One model suggests that the cholinergic pause is generated by intrinsic properties of CINs. When CINs come out of the early glutamatergic excitation, voltage-gated potassium channels (Kv7.2/7.3) open and induce an after-hyperpolarization that induces the pause. In this model DA plays a role in augmenting the intrinsically induced pause (Zhang et al., 2018). Consistent with this, thalamo-striatal stimulation induced a pause that was shortened but not fully abolished by a D2R antagonist (Cover et al., 2019). However, in earlier influential slice physiology experiments, the pause induced by thalamic stimulation was fully blocked by D2R antagonism suggesting that activation of DA release from intrastriatal DA terminals was responsible for pause generation (Ding et al., 2010).

One limitation of the mechanistic studies in rodents has been that they relied on stimulation experiments rather than on DA evoked by natural stimuli. While the early NHP studies suggested necessity for DA in inducing the pause during behavior, these studies lacked the cellular specificity for excluding the possibility that the effects of pharmacological D2R blockage were due to inhibiting D2Rs on CINs vs other neuronal populations.

Here, we used genetically encoded biosensors (Labouesse & Patriarchi, 2021) to simultaneously monitor DA and ACh in the dorsal striatum during behavior in mice with pharmacological blockade and/or selective ablation of D2Rs from CINs. Using this approach, we addressed the question whether the natural stimulus induced pause is fully dependent on DA or not. We first determined whether changes in DA and ACh levels occur simultaneously to reward-predicting stimuli in mice as has been shown in NHPs via electrophysiological recordings of DA and TAN neurons (Morris et al., 2004). *In vivo* imaging of ACh and DA levels revealed cue-induced decreases in striatal ACh and increases in DA levels, confirming the ability to measure concomitant ACh dips and DA peaks with functional imaging. Using a Pavlovian learning task, we confirmed that both signals co-occur and develop in parallel during the training of the task. Using a simpler reinforcement task that enables better quantification of the neuromodulator signals, we quantified cue-induced changes in DA and ACh changes after manipulating D2R function. We found that selective ablation of D2Rs from CIN or blocking D2Rs in control mice with the selective D2R antagonist eticlopride did not abolish the stimulus-induced decrease in ACh levels. Rather it shortened the length of dip and enhanced ACh rebound levels in a dose dependent manner. This indicates that DA is necessary for controlling the overall shape of the ACh dip. During simultaneous recordings experiments, the relationship between DA and ACh was strongest in response to reward predicting cues but still present during the intertrial interval supporting a general mechanism by which DA coordinates ACh levels. At the behavioral level, D2R antagonism increased latency to lever press to a reward-paired lever, but this relationship was abolished when we inactivated CIN D2Rs. Moreover, cue evoked changes in ACh levels correlated with the latency to press, altogether supporting a role of the ACh dip in motivated behavior.

## Materials and methods

### Animals

Adult male and female C57BL/6J (JAX stock # 000664) mice (Figures 1–6 & 10) were bred in house. For control and KO animals (Figures 7–10): double-transgenic mice were generated by crossing heterozygous ChAT-ires-Cre mice (Rossi et al., 2011) (JAX stock #006410) to homozygous *Drd2^fl/fl^* (Drd2^fl/fl^) mice (Bello et al., 2011). Control (Drd2^fl/fl^) and *ChATDrd2KO* (ChAT-ires-Cre x Drd2^fl/fl^) mice are littermates, bred in house and back crossed onto C57BL/6J background. Mice were housed 1-4 per cage for most experiments on a 12-hr light/dark cycle, and all experiments were conducted in the light cycle. All experimental procedures were conducted following NIH guidelines and were approved by Institutional Animal Care and Use Committees by Columbia University and the New York State Psychiatric Institute.

### Pharmacology

Intraperitoneal injections of saline, eticlopride (Tocris Cat. No. 1847) (0.1, 0.25, 1.0, 2.5 and 5.0 mg/kg) or scopolamine (Tocris Cat. No. 1414) (15 mg/kg) were administered 1hr before behavioral testing. To generate a dose-response curve with eticlopride, saline days alternated with drug days and the drug was administered in order from the lowest to highest dose.

### Surgical procedures

Mice (≥ 8 weeks old) were induced with 4% isoflurane and maintained at 1-2% throughout the procedure. Mice were bilaterally injected with 450 nL/hemisphere with either AA5-hSYN-dLight1.2 (Addgene) (Patriarchi et al., 2018) or AA5-hSYN-ACh3.0 (Vigene) (Jing et al., 2020) (also known as GRAB-ACh3.0) into separate hemispheres of the dorsal medial striatum (DMS) using stereotactic Bregma-based coordinates: AP, +1.1mm; ML ±1.4 mm; DV, −3.1 mm, −3.0 mm and −2.9 mm (150 nL/DV site). For the optogenetic inhibition experiment, mice were co-injected unilaterally with AA5-hSYN-ACh3.0 and AAV5-EF1α-DIO-eNpHR3.0 (UNC Vector Core) into the DMS. Following virus injection, 400-μm fiber optic cannulas (Doric, Quebec, Canada) were carefully lowered to a depth of −3.0 mm and fixed in place to the skull with dental cement anchored to machine mini-screws. Groups of mice used for experiments were housed in a counterbalanced fashion that accounted for sex, age, and home cage origin. Cannula-implanted mice began behavioral training 4 weeks after surgery. At the end of experiments, animals were perfused, and brains were processed post-*hoc* to validate virus expression and optic fiber location as in (Gallo et al., 2021).

### In vivo fiber photometry and optogenetics

Fiber photometry equipment was set up using two 4-channel LED Driver (Doric) connected to two sets of a 405-nm LED and a 465-nm LED (Doric, cLED_405 and cLED_465). The 405 nm LEDs were passed through 405-410 nm bandpass filters, while the 465 nm LEDs were passed through a 460-490 nm GFP excitation filters using two 6-port Doric minicubes. 405 and 465 LEDs were then coupled to a dichroic mirror to split excitation and emission lights. Two low-autofluorescence patch cords (400 μm/0.48NA, Doric) arising from the 2 minicubes were attached to the cannulas on the mouse’s head and used to collect fluorescence emissions. These signals were filtered through 500-540 nm GFP emission filters via the same minicubes coupled to photodetectors (Doric, gain set to DC Low). Signals were sinusoidally modulated, using Synapse® software and RZ5P Multi I/O Processors (Tucker-Davis Technologies), at 210 Hz and 330 Hz (405 nm and 465 nm, respectively) to allow for low-pass filtering at 3 Hz via a lock-in amplification detector. 405 and 465 nm power at the patch cord were set to 30 μW or below. For acute optogenetic inhibition via eNpHR3.0, amber light (595 nm LED, Doric) was applied through the same optic fiber using a short and long optogenetic protocol: (i) 500 ms square pulses at 1mW + 1s ramp down; (ii) 500 ms + 3s ramp down. The 595 nm light was passed through a 580-680 F2 port (photodetector removed) of the same 6-port minicube (Pisansky et al., 2019). The optogenetic experiment was performed in the home cage.

### Photometry data processing

All photometry and behavioral data utilized custom in-house MATLAB analysis scripts. Photometry signals were analyzed as time-locked events aligned to the lever extension (CRF) or tone onset (Pavlovian) of each trial. The 405 nm channel was used to control for potential noise/movement artifacts and the 465 nm channel was used to detect the conformational modulation of either the GACh3.0 sensor by ACh or the dLight1.2 sensor by DA. Both demodulated signals were extracted as a 15 s window surrounding the event, which was denoted as time = 0. Both signals were down sampled by a factor of 10 using a moving window mean. The change in fluorescence, ΔF/F (%), was defined as (F-F0)/F0 × 100, where F represents the fluorescent signal (465 nm) at each time point. F0 was calculated by applying a least-squares linear fit to the 405 nm signal to align with the 465nm signal (Calipari et al., 2016). To normalize signals across animals and sessions, we calculated a local baseline fluorescence value for each trial using the average of the 5 s period preceding the event and subtracted that from the signal. The daily average GACh3.0 and dLight1.2 traces were calculated using session average traces from individual mice. ACh dip and DA peak amplitudes were calculated as the maximal change of the signal that was at least 1 or 2 STD below or above the local baseline, respectively. ACh dip duration was calculated using the last and the first zero crossings preceding and following the dip. Total AUC was calculated as the area of all three components of the ACh signal (initial peak, dip and rebound). Negative AUC was calculated as the area for only the negative component. Rebound AUC was calculated as the area of the positive component immediately following the dip. The AUC analysis was restricted to a 5 s time window following the task event. Individual CRF trial (ΔF/F (%)) traces were used for correlation analysis for CRF trials. For the ITI correlation, we examined any interaction between dLight1.2 and GACh3.0 regardless of event size during the variable 40 s of the ITI.

### Operant apparatus

Four operant chambers (model Env-307w; Med-Associates, St. Albans, VT) equipped with liquid dippers were used. Each chamber was in a light- and sound-attenuating cabinet equipped with an exhaust fan, which provided 72-dB background white noise in the chamber. The dimensions of the experimental chamber interior were 22 × 18 × 13 cm, with flooring consisting of metal rods placed 0.87 cm apart. A feeder trough was centered on one wall of the chamber. An infrared photocell detector was used to record head entries into the trough. Raising of the dipper inside the trough delivered a drop of evaporated milk reward. A retractable lever was mounted on the same wall as the feeder trough, 5 cm away. A house light located on the wall opposite to trough illuminated the chamber throughout all sessions.

### Dipper training

Four weeks after surgery, mice underwent operant training. Mice were weighed daily and food-restricted to 85–90% of baseline weight; water was available ad libitum. In the first training session, 20 dipper presentations were separated by a variable inter-trial interval (ITI) and ended after 20 rewards were earned or after 30 min had elapsed, whichever occurred first. Criterion consisted of the mouse making head entries during 20 dipper presentations in one session. In the second training session, criterion was achieved when mice made head entries during 30 of 30 dipper presentations.

### Pavlovian conditioning

Mice were trained for 16 consecutive days in a Pavlovian conditioning paradigm, which consisted of 12 conditioned stimulus-positive (CS+) trials and 12 unconditioned stimulus (CS-) trials occurring in a pseudorandom order. Each trial consisted of an 80-dB auditory cue presentation for 10 s of an 8 kHz tone or white noise (counterbalanced between mice) and after cue offset a milk reward was delivered only in CS+ trials, whereas no reward was delivered in CS-trials. There was a 100 s variable intertrial interval, drawn from an exponential distribution of times. Head entries in the food port were recorded throughout the session, and anticipatory head entries during the presentation of the cue were considered the conditioned response. Anticipatory responding was calculated as the difference in nose poking during the CS+ quintile with the maximum response (Q4 or 5) and the first quintile.

### Continuous reinforcement schedule (CRF)

For lever press training, lever presses were reinforced on a continuous reinforcement (CRF) schedule. Levers were retracted after each reinforcer and were presented again after a variable ITI (average 40 s). The reward consisted of raising the dipper for 5 s. The session ended when the mouse earned 60 reinforcements, or one hour elapsed, whichever occurred first. Sessions were repeated daily until mice achieved 60 reinforcements.

### Data analysis

Sample sizes were determined by performing statistical power analyses based on effect sizes observed in preliminary data or on similar work in the literature. Statistical analyses were performed using GraphPad Prism 9 (GraphPad), MATLAB (MathWorks). Data are generally expressed as mean ± standard error of the mean (SEM). Paired and unpaired two-tailed Student’s t-tests were used to compare 2-group data, as appropriate. Multiple comparisons were evaluated by one- or two-way ANOVA and Bonferroni’s post hoc test, when appropriate. In rare cases of values missing in repeated measures samples, the data were analyzed by fitting a mixed effects model, as implemented by Prism 9. Photometry correlation analyses were performed using Pearson’s correlation coefficients. A p-value of < 0.05 was considered statistically significant. Behavioral findings were replicated with mice from different litters, ages, or sexes. Investigators were blinded to the genotype of mice during behavioral assays as well as throughout the data analysis. Computer code for data analysis is available on Github (username: kmartyniuk1). Files are titled, “DA-ACh_Dualimaging_CRF” and “DA-ACh_Dualimaging_Pavlovian”.

## Results

### GACh3.0 allows for measuring fast decreases in task evoked ACh levels

First, we validated our experimental approach to simultaneously image changes in ACh and DA levels within the same mouse (Figure 1A) during an instrumental task, continuous reinforcement (CRF) (Figure 1B). We aligned our photometry signals to the lever extension, which with training becomes a reward predicting cue. After 3 days of training, we observed an increase in DA (red) and a decrease in ACh (blue) at lever extension presentation (Figure 1C). To confirm that the fluorescent indicator, GACh3.0, is measuring changes in ACh levels (and not movement artefacts or electrical noise), we measured the GACh3.0 signal in the presence of 15 mg/kg scopolamine, a M3R muscarinic antagonist, which targets the GACh3.0 parent receptor (M3R). We found that scopolamine, abolished an early increase and the following decrease in ACh levels in response to lever extension when compared to saline injections (Figure 1D).

**Figure 1.**
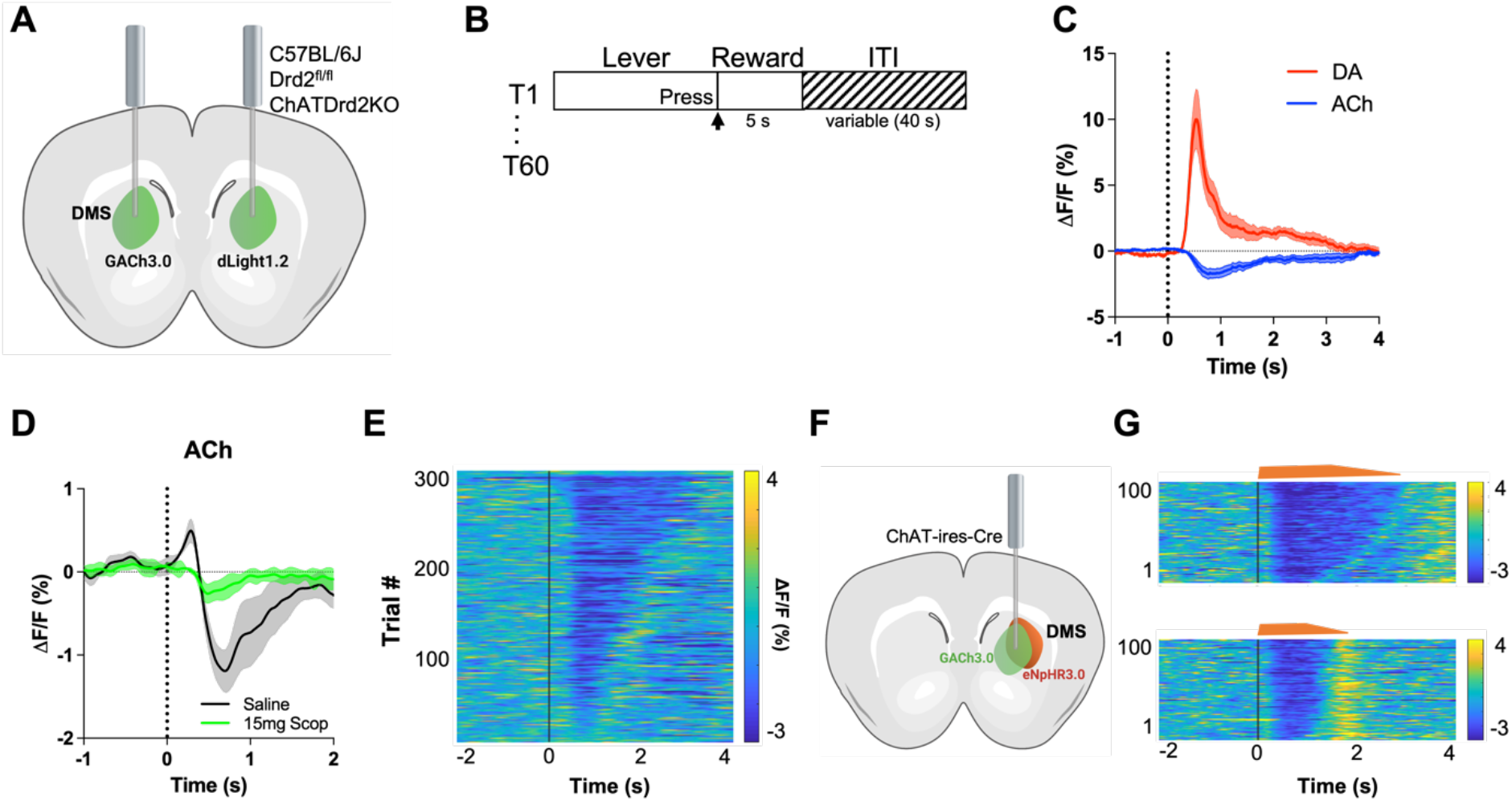
GACh3.0 reliably measures fast decreases in ACh during an instrumental task. **(A)** Schematic of the surgery setup. All mice were injected with both biosensor viruses (GACh3.0 and dLight1.2) in separate hemispheres of the DMS and counterbalanced across mice. Fiber photometry lenses were bilaterally implanted at the site of viral injection to simultaneously monitor ACh and DA in the same mouse. **(B)** CRF task design. Mice were trained to press a lever to retrieve a milk reward for 60 trials/day with a variable ITI (40 s). (**C)** Changes in fluorescence (ΔF/F) aligned to lever extension (timepoint= 0 s). DA levels (red) increased and ACh levels (blue) decreased, N= 5 mice in trained mice. **(D)** 15 mg/kg of scopolamine (green), a mAChR antagonist, blunts the initial ACh peak and dip compared to saline (black) confirming that the GACh3.0 sensor is reporting changes in ACh levels. N= 4 mice. **(E)** Heatmap of ACh responses aligned to lever extension (Time= 0 s) and sorted by length of ACh decrease for 300 individual trials (60 trials in 5 mice). **(F)** Schematic of the surgery setup. ChAT-ires-Cre mice were co-injected with GACh3.0 and Cre-dependent halorhodopsin into the DMS and a fiber photometry lens was implanted at the site of viral injection. **(G)** Approximation of trials with short dips (bottom) and long dips (top) using the short and long optogenetic inhibition protocol (100 trials, 20 trials/ 5 mice).

To confirm that the GACh3.0 sensor has the kinetics to measure a rapid decrease in ACh levels, we expressed the inhibitory opsin eNpHR3.0 in ChAT-ires-Cre mice to selectively inhibit CINs. Lever extension induced decreases in ACh levels within 250 ms showed varying lengths trial by trial (Figure 1E). Light activation of eNpHR3.0 induced a dip with even shorter latency (latency to dip onset 206.4 [186.8-226.1] ms, n=5 mice), which was followed by a rebound in ACh levels (Figure 1F). The optogenetic experiment was performed in the home cage. This is consistent with CINs displaying rebound activity after injecting hyperpolarizing currents in brain slices (Wilson, 2005) and optogenetic inhibition *in vivo* (English et al., 2012). Combined, these data show that GACh3.0 can measure fast decreases in ACh levels. It also indicates that ACh release and degradation and/or diffusion are tightly controlled by CIN neuron activity.

### Simultaneous development of DA and ACh signals in response to a reward predicting stimulus

To determine whether changes in DA and ACh levels in response to reward predicting stimuli are co-incident, we measured the release of DA and ACh during a classic Pavlovian reward learning task (Figure 2B). Using fiber photometry and genetically encoded fluorescent indicators, we simultaneously imaged DA and ACh in separate hemispheres within the same animal (Figure 2A). On Day 1 of training, we observed an increase in DA (red) and a decrease in ACh (blue) during unexpected reward following the offset of the CS+. Over training, we saw these changes in both DA and ACh shift to the onset of the CS+ tone, while decreasing to the now expected reward. We did not observe these changes during CS- trials. We then related the changes in DA and ACh to changes in anticipatory nose poking during the CS+ as a measure of learning. We found that both DA and ACh signals correlated well with anticipatory head poking in one animal (Figure 2C). However, other mice did not show any anticipatory responding as this task is non-contingent and head poking is not required to obtain the reward during CS+ trials. These findings indicate that DA and ACh signals co-develop with learning in response to a reward predicting stimulus.

**Figure 2.**
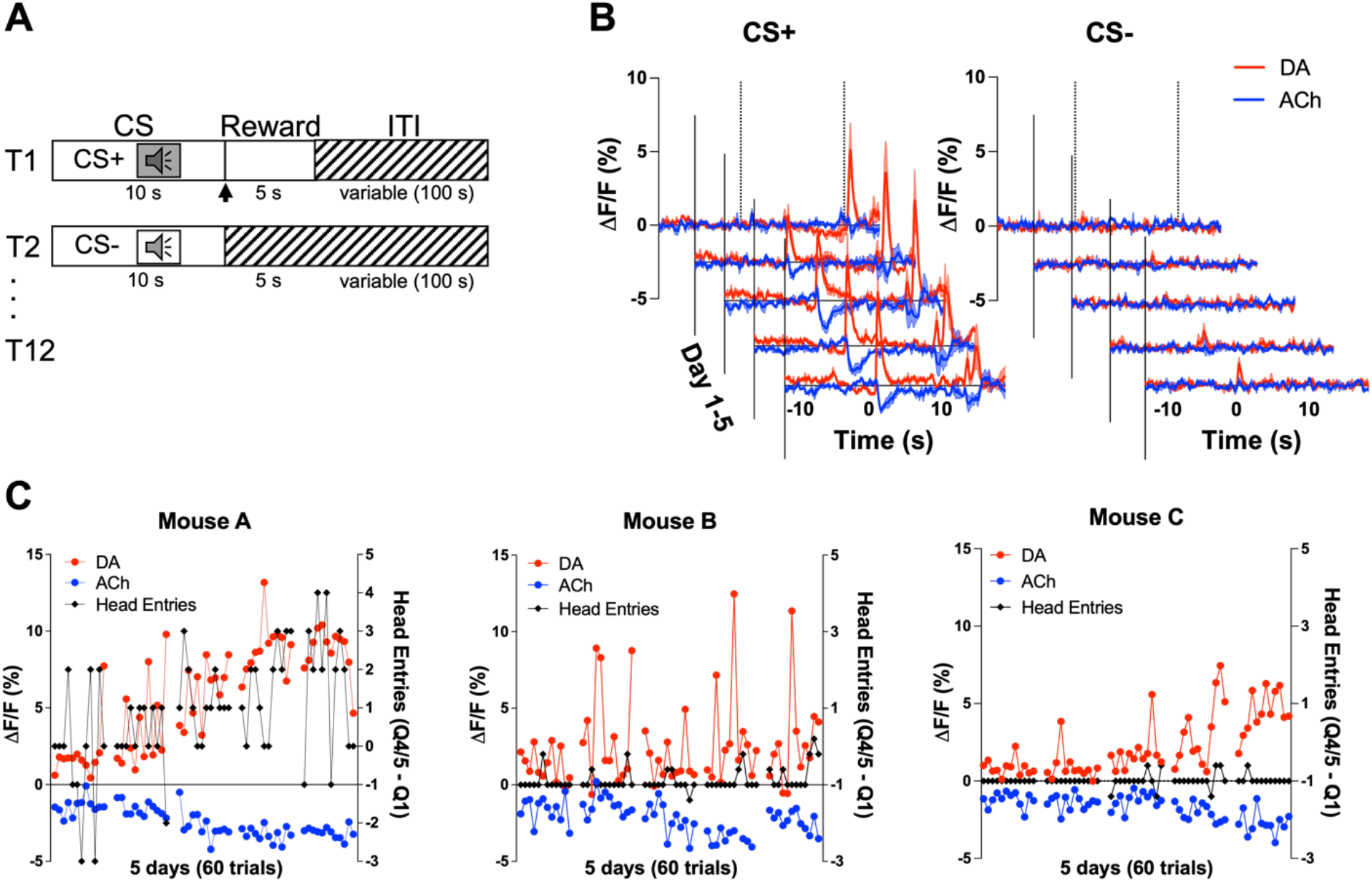
Co-development of DA and ACh signals to a reward predicting cue. **(A)** Pavlovian task design. Mice were trained on 24 (12 CS+, 12 CS-) trials/day for 5 days. Each trial starts with a 10 s tone (CS+ or CS-). At the end of the CS+ a dipper comes up presenting a milk food reward for 5 s. There is an intertrial interval (ITI) variable in length (100 s). **(B)** Changes in fluorescence (ΔF/F (%)) over 5 days of training for DA (red) and ACh (blue) aligned to CS+ (left) and CS- (right) onset. Signals were averaged over 12 CS+ and 12 CS- trials/day, N= 3 mice. **(C)** Maximum change in DA peak (blue) and ACh dip (red) after CS+ onset over 5 days of training (60 trials) for Mouse A (left). Anticipatory responding (black) is calculated as the difference in nose poking during the CS+ quintile with the maximum responses (Q4 or 5) and the first quintile. Correlations between DA and ACh maxima and behavioral responding: r= 0.4, p<0.002 and r= −0.41, p<0.002, respectively. We did not observe the same correlation between DA/ACh and anticipatory responses in Mouse B (middle) or Mouse C (right).

### D2 receptor blockade dose dependently decreases ACh dip lengths and enhances the rebound in ACh levels

To determine if the cue induced ACh dip is dependent on DA activation of D2Rs, we used the CRF task as it allows for more trials per session aiding the quantification of the signal. After systemic delivery of the D2R antagonist eticlopride we found a dose-dependent shortening of the ACh dip, which uncovered a rebound following the pause (Figure 3A). We quantified these changes by calculating the area under the curve (AUC), dip length and dip amplitude. We found that eticlopride significantly reduced the negative AUC (Figure 3B), increased the rebound AUC (Figure 3C), increased the total AUC (Figure 3D), and decreased the dip length (Figure 3E), while the dip amplitude was not affected by D2R antagonism (Figures 3F). This suggests that D2Rs do not participate in the initial induction of the ACh dip but do increase the length of the dip and prevent rebound activity following the ACh dip. Since DA neurons are inhibited by D2 auto-receptors, we also analyzed the effect of D2R antagonism on cue induced DA release and quantified changes in peak amplitude and AUC (Figure S1A). We found an overall effect of drug increasing the peak amplitude (Figure S1B) with the most prominent increase between saline and 0.1 mg/kg. There was no overall effect of drug on the AUC (Figure S1C). These results confirm that blocking D2 auto-receptors on DA neurons increases phasic DA release.

**Figure 3.**
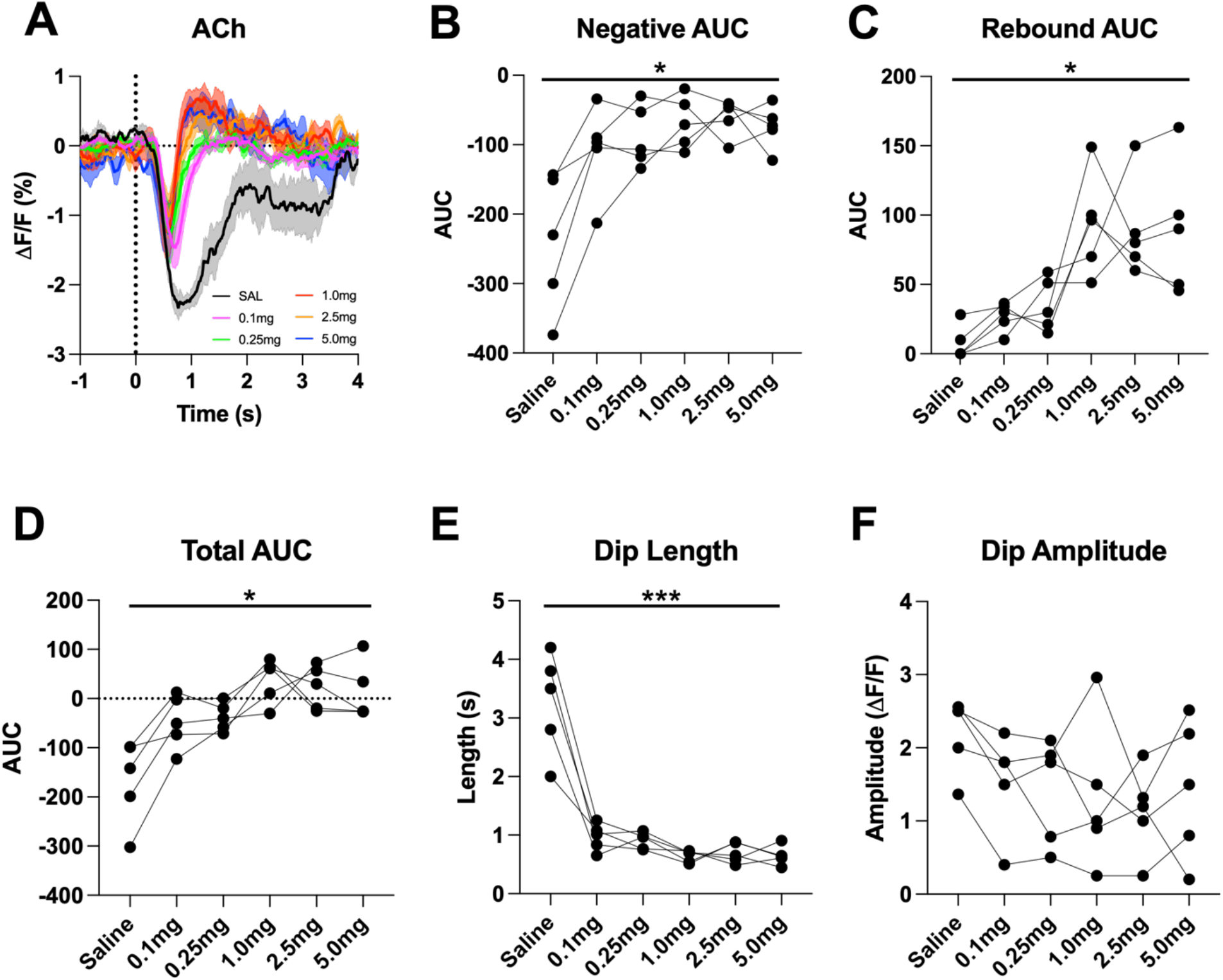
D2R antagonism decreases ACh dip length and enhances rebound. **(A)** Changes in ACh fluorescence (ΔF/F (%)) aligned to lever extension with saline (black) and increasing doses of eticlopride: 0.1 mg/kg (pink), 0.25 mg/kg (green), 1.0 mg/kg (red), 2.5 mg/kg (orange) and 5.0 mg/kg (blue). **(B)** Negative AUC is reduced by eticlopride in a dose-dependent manner (F_(1.694, 6.777)_ = 8.756, p = 0.0150). **(C)** The rebound AUC is increased by eticlopride in a dose-dependent manner (F_(1.549, 6.197)_ = 8.833, p= 0.0181) **(D)** Total AUC is increased by eticlopride in a dosedependent manner (F_(1.612, 6.448)_ = 8.724, p = 0.0170). **(E)** Dip length is decreased by eticlopride in a dose-dependent manner (F_(1.392, 5.569)_ = 36.37, p = 0.0009). **(F)** The dip amplitude was not affected by eticlopride (F_(2.063, 8.251)_ = 1.864, p = 0.2147).

Individual CRF trials revealed varying lengths of lever-extension aligned ACh dips that we sorted by lever press latency using a heatmap (Figure 4A). Based on this heatmap, we observed longer dips associated with quick press latencies and two smaller dips with slower press latencies with the second dip co-occurring with the lever press. Thus, for press latencies < 2 s the ACh is a combination of a cue induced and movement associated pause. To separate the cue induced pause from the movement induced pause, we analyzed trials with press latencies > 2 s. We still observed a decrease in the ACh dip duration with increasing doses of eticlopride (Figure 4B). Quantification of the negative AUC revealed a non-significant but trending decrease with increasing doses of eticlopride (Figure 4C), while the rebound AUC increased (Figure 4D), the total AUC increased (Figure 4E) and the dip length (Figures 4F) decreased. Eticlopride had no effect on the ACh dip amplitude (Figure 4G). We also examined the effect of D2R antagonism on cue induced DA release for trials with press latencies > 2s (Figure S2A). Quantification of DA peak amplitude (Figure S2B) and AUC (Figure S2C) revealed an overall increase in both measures. Moreover, we found a significant increase between saline and 0.1 mg/kg eticlopride for DA peak amplitude (Figure S2B). Taken together, these results demonstrate that the cue induced ACh dip and rebound levels are regulated by cholinergic D2Rs.

**Figure 4.**
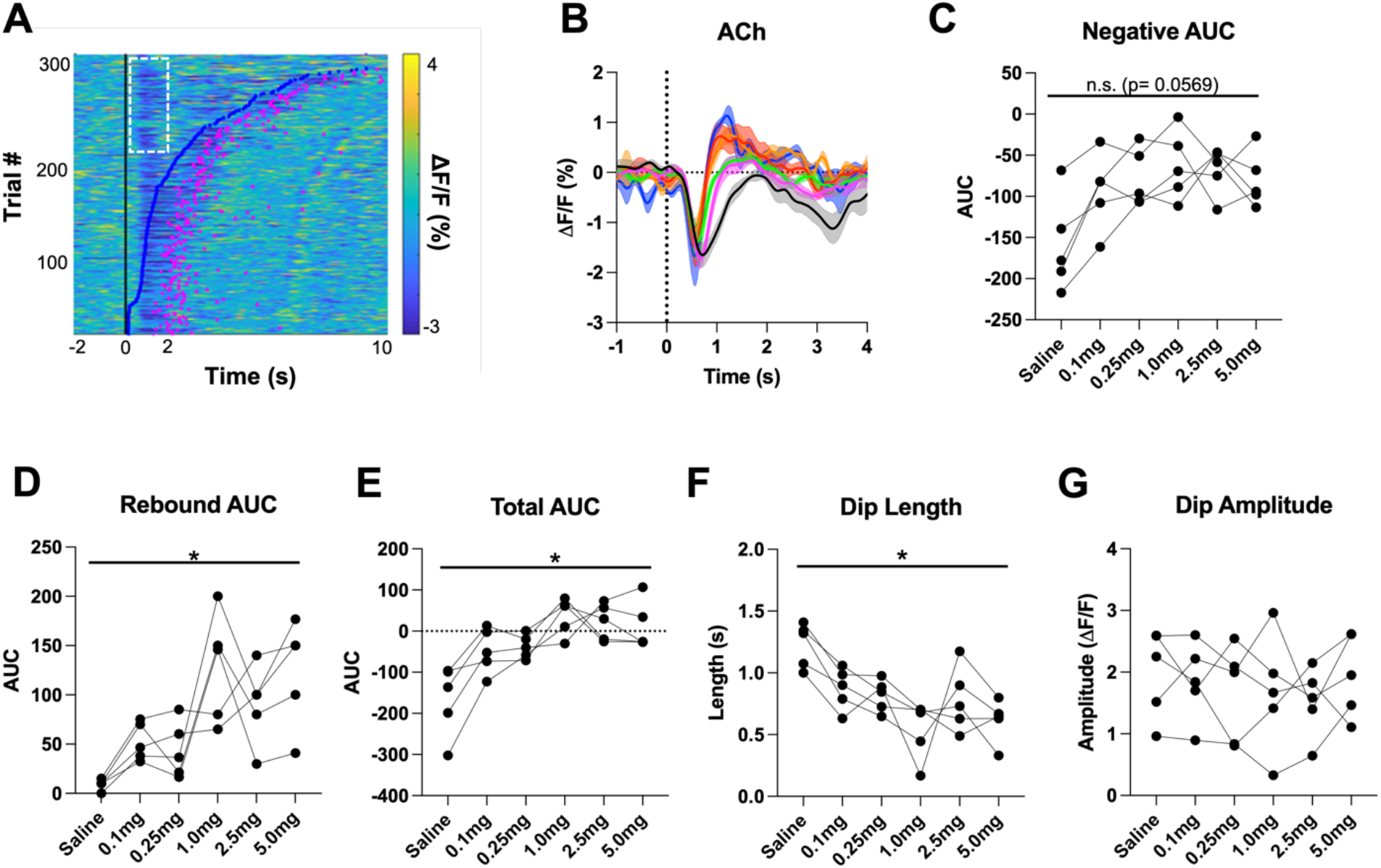
D2R antagonism shortens cue evoked ACh dip and enhances rebound. **(A) H**eatmap of ACh responses aligned to lever extension (Time= 0 s) for 300 individual trials (60 trials in 5 mice) and sorted by response length (bottom). Blue dots show the lever press, and the pink dots show the head entry for each trial. White dashed box represents the cue evoked ACh response to the lever extension where press latencies > 2 s. **(B)** Changes in ACh fluorescence (ΔF/F (%)) aligned to lever extension for only trials with press latencies > 2 s with increasing doses of eticlopride. **(C)** Negative AUC is reduced by eticlopride in a dose-dependent manner (F_(2.237, 8.950)_ = 3.911, p = 0.0569). **(D)** Rebound AUC is enhanced by eticlopride in a dose-dependent manner (F_(1.667, 6.668)_ = 8.143, p = 0.0184). **(E)** Total AUC was increased by eticlopride in a dose-dependent manner (F_(1.597, 6.387)_ = 8.542, p = 0.0182). **(F)** Dip length was significantly decreased by eticlopride in a dose-dependent manner (F_(1.657, 6.628)_ = 6.729, p = 0.0284). **(G)** Eticlopride had no effect on the dip amplitude (F_(2.722, 10.89)_ = 0.5379, p = 0.6503).

### D2R blockade decreases negative and enhances positive correlations between DA and ACh

We further determined the relationship between ACh and DA levels within trials using a Pearson’s correlation analysis. Using a lag analysis, we temporally shifted the ACh recording behind or in front of the DA recording to identify maximal points of correlation. During CRF trials, the strongest correlation is a negative correlation (Figure 5A, label 1, saline: Pearson’s r= −0.475 ± 0.037, N=5) that occurs when ACh lags DA (Lag= −178.92 ms ± 14.38 ms). This negative correlation, which reflects the ACh dip that follows the DA peak, is reduced with eticlopride in a dose-dependent manner (Figure 5). Next, we found a small positive correlation (Figure 5A, label 2, saline: Pearson’s r= 0.039 ± 0.014) when ACh lags DA (Lag= −1.5 s ± 0.138 s). This positive correlation, which reflects the rebound in ACh, is significantly increased with eticlopride (Figure 5C).

**Figure 5.**
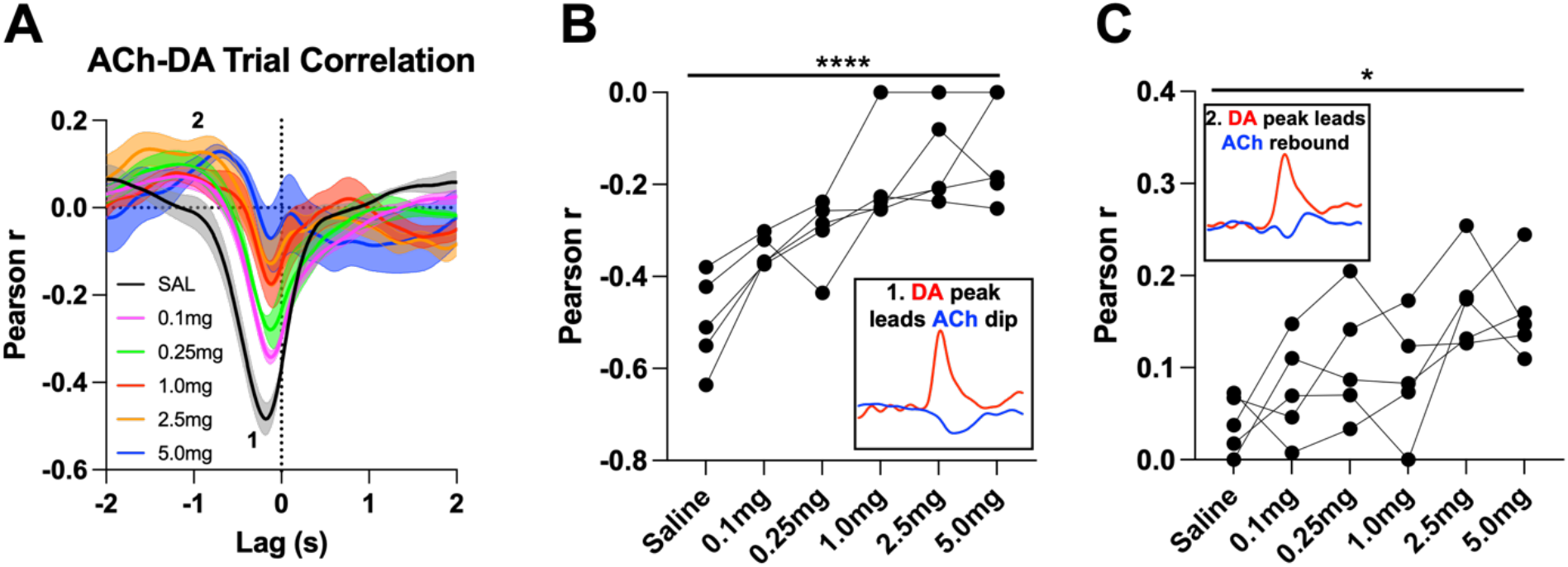
Task dependent ACh-DA interactions are altered by D2R antagonism at lever extension. **(A)** Correlation between ACh and DA during CRF trials with increasing doses of eticlopride in 5 C57BL/6J mice. The ACh signal moved in front of or behind the DA signal to identify points of highest correlation. The first correlation is a negative correlation (1) with ACh lagging DA (Lag= −178.92 ms ± 14.38 ms) and the second correlation is a positive correlation (2) with ACh lagging DA (Lag= −1.5 s ± 0.138 s). **(B)** The negative correlation with the DA peak leading the ACh dip (inset) is significantly reduced dose-dependently by eticlopride (F_(3.082, 12.33)_ = 18.67, p < 0.0001) **(C)** The positive correlation with the DA peak leading the ACh rebound (inset) is enhanced by eticlopride in a dose-dependent manner (F_(2.325, 9.299)_ = 4.731, p= 0.0346).

We then analyzed these correlations during the intertrial interval (ITI) to determine whether they are only present during stimulus induced DA/ACh signals or may represent a more general mechanism or coordination (Figure 6A). Of note, we looked for any interaction between DA and ACh regardless of event size. Like CRF trials, we observed two correlations during the ITI; DA peak leads ACh dip (Pearson’s r= −0.355 ± 0.065 and Lag= −212.34 ms ± 16.91 ms) and DA peak leads ACh peak/rebound (Pearson’s r= 0.058 ± 0.021 and Lag= −1.41 s ± 0.19 s). We found that eticlopride decreases the negative correlation in a dose-dependent manner (Figure 6B). Eticlopride also increased the positive correlation, which represents the ACh rebound (Figure 6C). These results indicate that DA-ACh correlations are dependent on D2Rs. While they are strong during salient cue presentations the relationship between both signals still exists during the intertrial interval reflecting a general mechanism of co-regulation.

**Figure 6.**
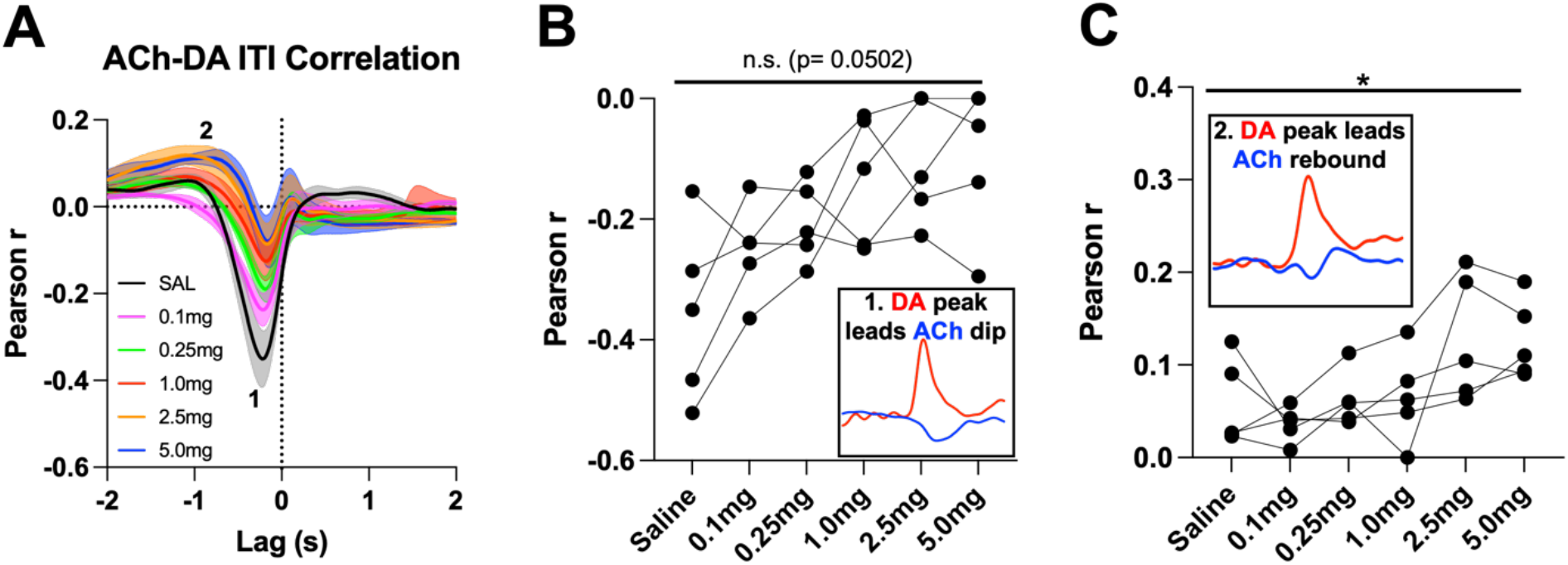
General ACh-DA interactions are altered by D2R antagonism during the ITI. **(A)** Correlation between ACh and DA during the ITI with increasing doses of eticlopride in C57BL/6J mice. We observe the same two correlations during the ITI: a negative correlation (1) with ACh lagging DA (Lag= −212.34 ms ± 16.91 ms) and a positive correlation (2) with ACh lagging DA (Lag= −1.41 s ± 0.19 s) **(B)** The negative correlation with the DA peak leading the ACh dip (inset) is decreased by eticlopride in a dose-dependent manner (F_(1.900, 7.598)_ = 4.606, p= 0.0502). **(C)** The positive correlation with the DA peak leading the ACh rebound (inset) is increased dose-dependently by eticlopride (F_(2.118, 8.474)_ = 4.873, p= 0.0377).

### Genetic inactivation of D2Rs from CINs decreases dip length

Systemic eticlopride injections block all D2Rs. To determine the specific modulatory role that D2Rs present in CINs play in the cholinergic pause, we used mouse genetics to selectively inactivate D2Rs from CINs (ChATDrd*2*KO mice). We observed a smaller and shorter ACh dip in ChATDrd2KO mice (Dip amplitude= 0.797ΔF/F ± 0.131 and Dip length= 0.931 s ± 0.105 s) compared to control mice (Dip amplitude= 1.50ΔF/F ± 0.141 and Dip length= 2.80 s ± 0.682 s) in trials with press latencies > 2 s (Figure 7A). Both, the dip amplitude, and dip length were significantly reduced between the two groups (Figure 7B-C). Note that the effects of D2R deletion differed from the highest dose of eticlopride in that ChATDrd2KO mice showed differences in the dip amplitude while eticlopride did not.

**Figure 7.**
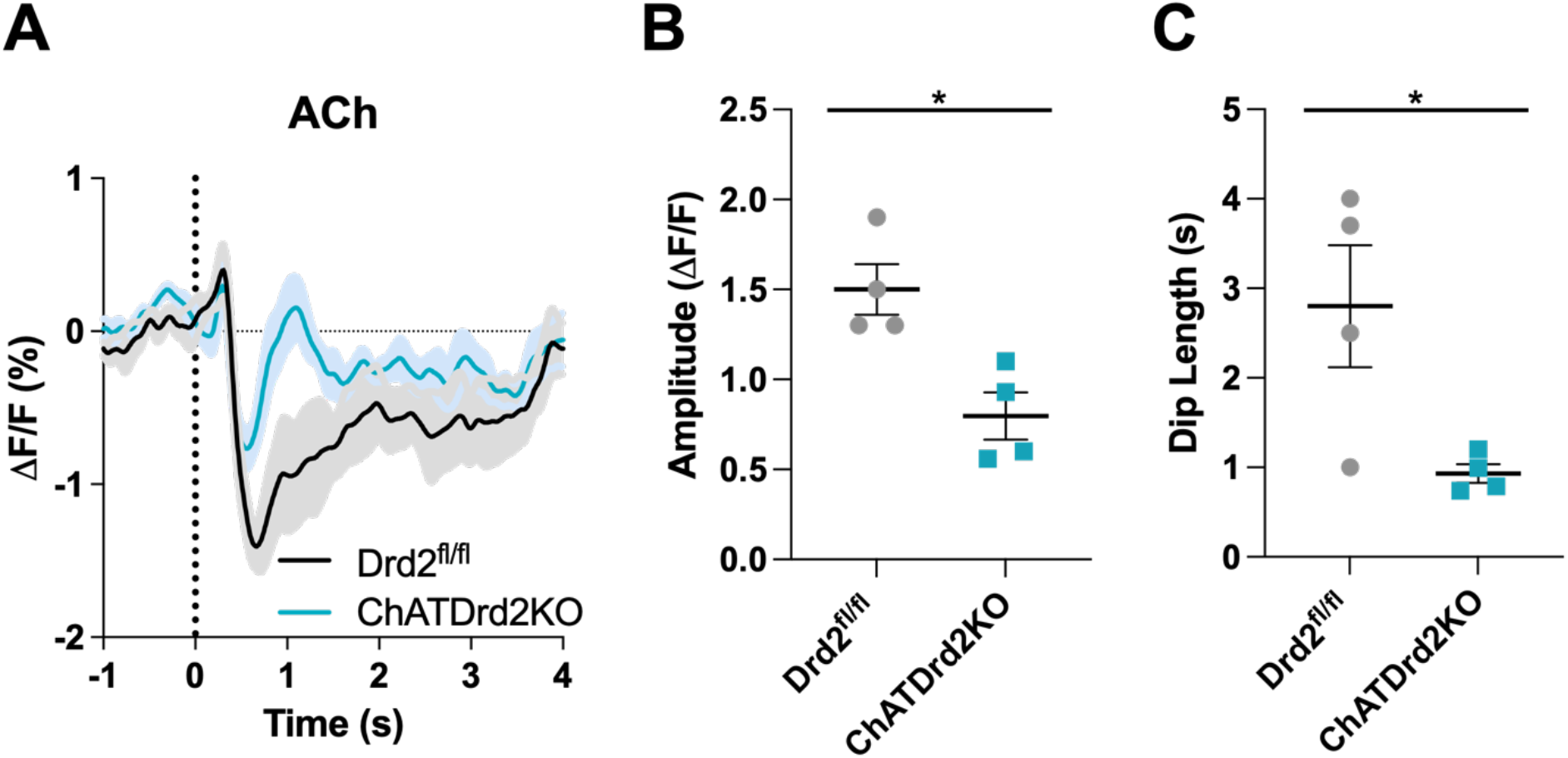
Selective D2R ablation from CINs alters the cue evoked ACh dip. **(A)** Changes in ACh fluorescence (ΔF/F (%)) aligned to lever extension for only trials with press latencies >2 s for Drd2^fl/fl^ control (black) and ChATDrd2KO (blue) mice, N= 4/genotype. **(B)** Dip amplitude is significantly smaller in ChATDrd2KO animals compared to controls (p= 0.0107). **(C)** Dip length is significantly shorter in ChATDrd2KO mice compared to Drd2^fl/fl^ controls (p= 0.0351).

In contrast to ACh levels, stimulus induced DA release was not altered in ChATDrd2KO mice (Figure S3). This result indicates that loss of cholinergic D2Rs does not affect stimulus induced DA release and confirm that the effects of DA regulation of the ACh dip are mediated by CIN D2Rs and not an indirect effect by potential changes in DA levels.

### DA-mediated changes in ACh levels are dependent on CIN D2Rs

Next, we determined if D2Rs present in CINs are necessary for the effect of D2R antagonism on modulating the ACh dip. Control Drd2^fl/fl^ mice were more sensitive to eticlopride than the C57BL/6J wild-type mice of Figure 4 as they did not complete any trials with the two highest doses, 2.5 mg/kg, and 5.0 mg/kg (Figure 8A-F). Quantification of the ACh dip using the 3 lower doses revealed a decrease in the negative AUC (Figure 8B), an increase in the rebound AUC (Figure 8C), an increase in the total AUC (Figure 8D) and a decrease in dip length (Figure 8E) that were comparable to what we measured in the C57BL/6J mice (Figure 4). Like the C57BL/6J mice, there was no effect on ACh dip amplitude with eticlopride (Figure 8F). In contrast, in ChATDrd2KO mice, we observed no change in the ACh dip with eticlopride (Figure 8G) and there was no effect of eticlopride on the negative AUC (Figure 8H), the rebound AUC (Figure 8I), total AUC (Figure 8J), dip length (Figure 8K), or dip amplitude (Figures 8L). These results confirm that D2Rs present in CINs are key players in the modulation of the ACh signal elicited by D2R antagonism.

**Figure 8.**
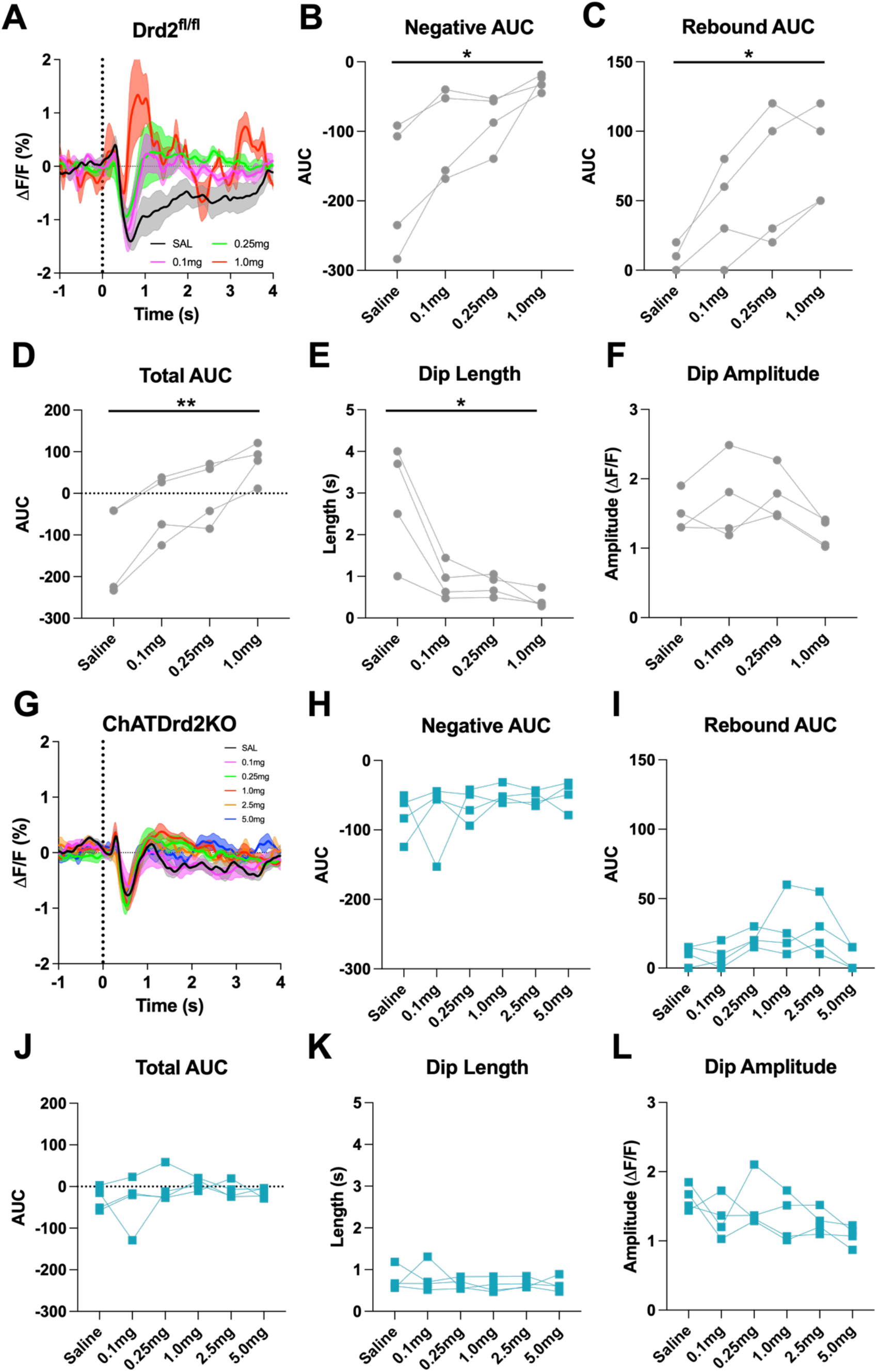
D2R antagonism does not alter the cue evoked ACh dip in ChATDrd2KO mice. **(A)** Changes in ACh fluorescence (ΔF/F (%)) aligned to lever extension for only trials with press latencies > 2 s for Drd2^fl/fl^ control mice with increasing doses of eticlopride. **(B)** Negative AUC is decreased by eticlopride in a dose-dependent manner (F_(1.387, 4.160)_ = 8.541, p = 0.0381). **(C)** Rebound AUC is increased by eticlopride in a dose-dependent manner (F_(1.642, 4.925)_ = 10.21, p = 0.0195). **(D)** Total AUC is increased dose-dependently by eticlopride (F_(1.525, 4.676)_ = 23.14, p = 0.0047). **(E)** Dip length is decreased by eticlopride in a dose-dependent manner (F_(1.664, 4.992_) = 9.279, p = 0.0226). **(F)** Dip amplitude is not affected by eticlopride (F_(1.433, 4.300)_ = 6.056, p = 0.0606). **(G)** Changes in ACh fluorescence (ΔF/F (%)) aligned to lever extension for only trials with press latencies > 2 s for ChATDrd2KO mice with increasing doses of eticlopride. **(H)** Negative AUC is not affected by eticlopride (F_(1.663, 4.990)_ = 0.7919, p = 0.4803). **(I)** Rebound AUC is not affected by eticlopride (F_(1.706, 5.119)_ = 2.857, p = 0.1484). **(J)** Total AUC is not affected by eticlopride (F_(1.844, 5.532)_ = 1.079, p = 0.3958). **(K)** Dip length is not affected by eticlopride (F_(1.848, 5.545)_ = 0.4380, p = 0.6516). **(L)** Dip amplitude is not affected by eticlopride (F_(2.073, 6.219)_ = 2.546, p = 0.1551).

### DA-mediated changes in DA-ACh correlations are dependent on CIN D2Rs

Next, we assessed the effect of CIN D2Rs in the ACh-DA coregulation, again using Pearson’s r correlation analysis and lag analysis. We found that the interaction between DA and ACh was greatly reduced (> 2-fold) in ChATDrd2KO mice compared to Drd2^fl/fl^ controls (Figure 9). The negative correlation with ACh lagging DA was significantly smaller in ChATDrd2KO mice during both CRF trials (Figure 9A-B) and the ITI (Figure 9C-D) compared to Drd2^fl/fl^ controls. We further examined the role of CIN D2Rs in the synchronization of DA and ACh activity using eticlopride to transiently block CIN D2Rs. Drd2^fl/fl^ control mice showed a strong negative correlation during CRF trials (Figure 9A: Pearson’s r= −0.521 ± 0.038, N=4) with ACh lagging DA (Lag= −167.12ms ±18.82 ms) that was reduced by eticlopride (Figure S4B). In addition, a rebound in ACh activity was revealed, which is measured as a positive correlation (Figure S4A: Pearson’s r= 0.050 ± 0.030, N=4) with ACh lagging DA (Lag= −1.51 s ± 0.138 s) that was increased with increasing doses of eticlopride (Figure S4C). During the ITI, the negative correlation between ACh and DA with ACh lagging DA (Figure S5A: Lag= −213.81 ms ± 34.14 ms) was attenuated by eticlopride (Figure S5B), while the positive correlation with ACh lagging DA (Lag= −1.39 s ± 0.208 s) was not significantly affected by eticlopride (Figure S5C). In ChATDrd2KO mice, neither the negative correlation between ACh and DA (Figure 9A: Pearson’s r= −0.231 ± 0.046, N=4) with ACh lagging after DA (Lag= −179.41 ms ± 30.66 ms, N=4) nor the positive correlation with ACh lagging DA (Lag= −1.635 s ± 0.174 s) were affected by eticlopride (Figure S6C). During the ITI, we observed a smaller negative correlation between ACh and DA (Figure 9C: Pearson’s r= −0.153 ± 0.034, N=4) with ACh lagging DA (Lag= −282.62 ms ± 93.81 ms) that was significantly smaller in ChATDrd2KO mice compared to control mice (Figure 9D) and was not affected by eticlopride (Figure S7A-B). These results it is that D2Rs in CINs that regulate ACh dip and rebound levels.

**Figure 9.**
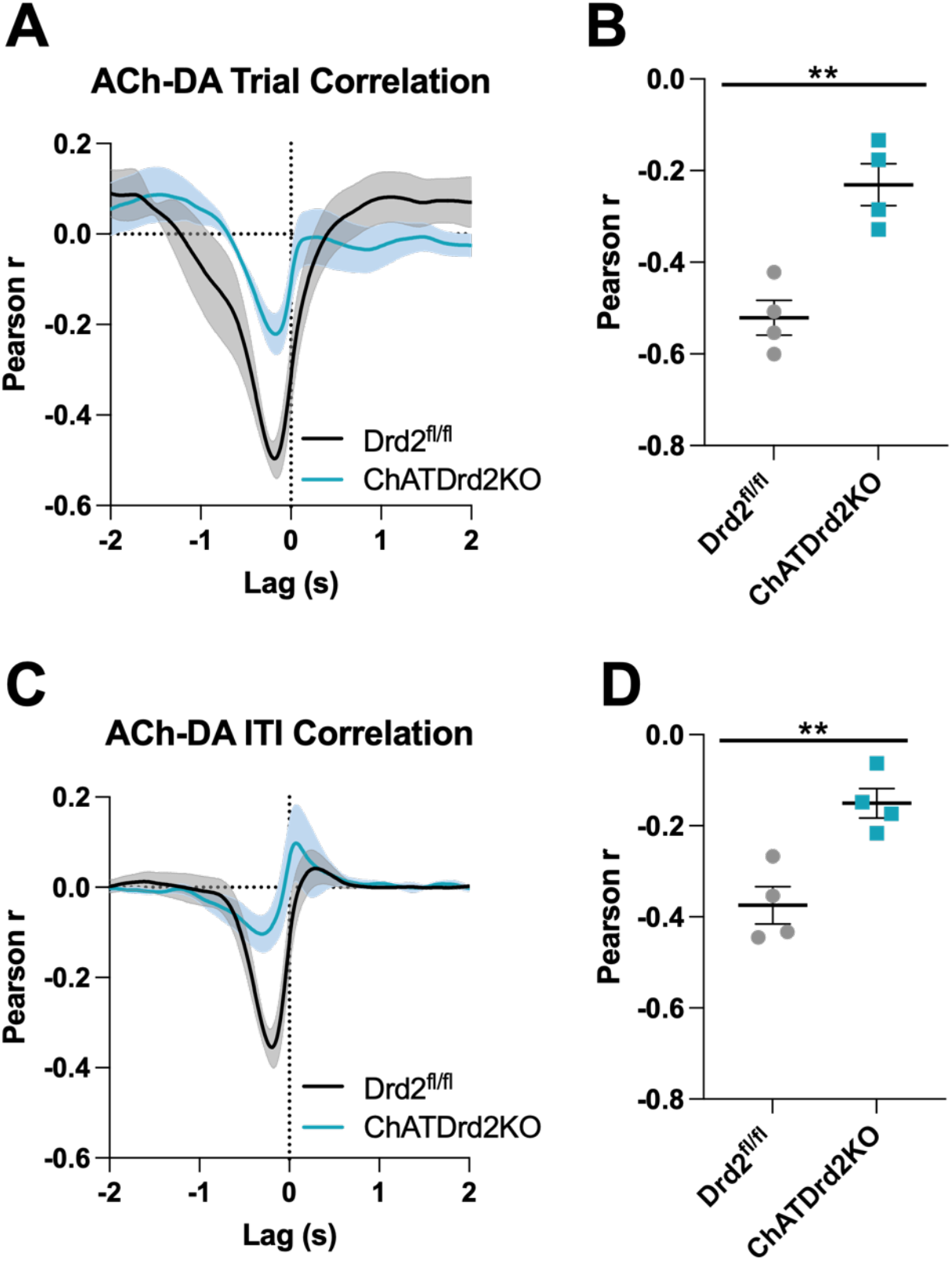
ACh-DA interactions are reduced in ChATDrd2KO mice. **(A)** Task evoked correlation between ACh and DA for Drd2^fl/fl^ control (black) and ChATDrd2KO (blue) mice. **(B)** The negative correlation with ACh lagging DA is significantly reduced in ChATDrd2KO mice compared to Drd2^fl/fl^ controls (p= 0.0155). **(C)** Correlation between ACh and DA during the ITI for Drd2^fl/fl^ control (black) and ChATDrd2KO (blue) mice. **(D)** The negative correlation of ACh lagging DA is significantly reduced in ChATDrd2KO mice compared to Drd2^fl/fl^ controls (p= 0.0052).

### D2R antagonism increases the latency to press in a CIN D2R dependent manner

We then determined if manipulating CIN D2R function affects behavior in the CRF task. Since D2R blockade induces catalepsy (Kharkwal et al., 2016) we wondered whether Drd2 ablation or D2R antagonism alters behavioral responding (latency to press in the task), an indicator of motivated behavior. In C57BL/6J mice, we found that eticlopride significantly increased lever press latency in a dose-dependent manner (Figures 10A). Eticlopride had no effect on lever press latency in ChATDrd2KO mice, compared to Drd2^fl/fl^ control mice (Figure 10B). Next, we determined if the size of the stimulus induced ACh dip correlates with behavioral responding. To do this, we analyzed the correlation between the AUC and lever press latency for trials with press latencies > 2s to isolate the stimulus induced ACh dip from the lever press associated dip. In C57BL/6J mice of Figures 3–6, we found a positive correlation between total AUC and press latency (Figure 10C). Similarly, in Drd2^fl/fl^ control mice, we found a similar positive correlation between total AUC and press latency (Figure 10E). This correlation was mainly driven by the ACh dip as the negative AUC positively correlated with press latency for C57BL/6J (Figure 10D) and Drd2^fl/fl^ control (Figure 10F) mice. This positive correlation between total AUC and press latency was disrupted in ChATDrd2KO animals (Figure 10G), while there remained a weaker positive correlation between the negative AUC and press latency (Figure 10H). These data suggest that D2R mediated regulation of cholinergic ACh levels contribute to the regulation of motivated behavior.

**Figure 10.**
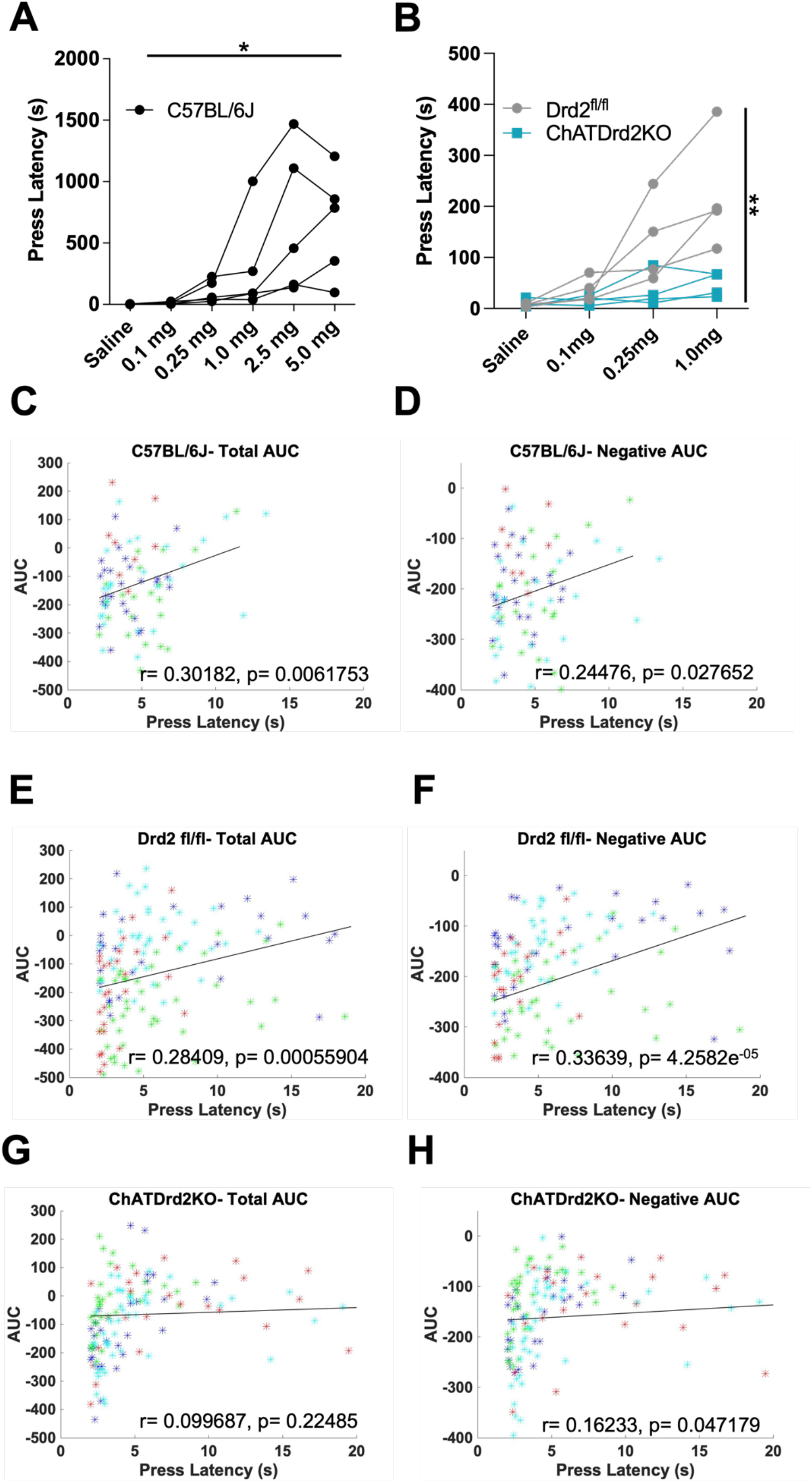
Behavioral responding correlates with ACh event size but is affected by D2R antagonism and ablation. **(A)** Lever press latency is increased by eticlopride in a dose-dependent manner in C57BL/6J mice (F_(1.383, 5.533)_ = 6.369, p = 0.0427). **(B)** D2R antagonism does not increase lever press latency in ChATDrd2KO mice (blue squares) compared to Drd2^fl/fl^ controls (gray circles) (F_(3,18)_ = 5.664, p = 0.0065, eticlopride x genotype). **(C)** Total AUC positively correlates with lever press latency in C57BL/6J mice (r= 0.30182, p= 0.0061753). **(D)** Negative AUC positively correlates with lever press latency in C57BL/6J mice (r= 0.24476, p= 0.027652). **(E)** Total AUC positively correlates with lever press latency in Drd2^fl/fl^ control mice (r= 0.28409, p= 0.00055904). **(F)** Negative AUC positively correlates with lever press latency in Drd2^fl/fl^ control mice (r= 0.33639, p= 4.2582e^−05^). **(G)** Total AUC does not correlate with lever press latency in ChATDrd2KO mice (r= 0.099687, p= 0.22485). **(H)** Negative AUC positively correlates with lever press latency in ChATDrd2KO mice (r= 0.16233, p= 0.047179).

## Discussion

Here, we investigated the mechanism by which striatal DA regulates cue induced changes in ACh levels during behavior. Understanding this mechanism is important because both neuromodulators coincidentally signal salient cues or outcomes during learning and motivated behavior and thus DA may regulate behavior via regulating ACh levels (Apicella et al., 1992). Moreover, it addresses the longstanding question of whether the ACh dip is fully dependent on striatal DA.

By simultaneously recording task-evoked DA and ACh levels in mice we made several observations: First, we observed that changes in striatal DA and ACh levels are induced by reward-predicting stimuli and the time locked signals develop in parallel with learning. Second, we found that pharmacological and genetic inactivation of D2Rs does not completely abolish the stimulus-induced dip in ACh, but it does shorten the dip and enhances rebound levels. Third, using correlational analysis, we found a relationship between DA and ACh that was strongest in response to lever extension as a reward predicting cue but still present during the inter-trial interval. This relationship was disrupted by D2R inactivation. Fourth, we found that D2R antagonism increased latency to lever press during behavior, but this was abolished when we inactivated CIN D2Rs. Lastly, the size of the cue evoked ACh dip but not rebound correlated with lever press latency, even for lever presses that happen long after the cued signal has ended. Altogether, these findings indicate that DA and cholinergic D2Rs are necessary for controlling the shape of the ACh dip and the coordinated activity between DA and ACh during reward driven behaviors. Moreover, the cue induced ACh dip correlates and therefore may drive behavioral responding during motivated behavior.

### Cue induced changes in striatal DA and ACh levels are time locked and develop in parallel with learning

The changes in DA and ACh levels that we recorded in the Pavlovian conditioning task are consistent with DA neurons and CINs encoding unexpected rewards and reward-predicting cues (Aosaki T, 1994b; Joshua et al., 2008; Morris et al., 2004; Schultz et al., 1997; Watanabe & Kimura, 1998). Like these previous studies, that assayed neuronal activity, we see a robust increase in DA levels and a decrease in ACh levels to unexpected reward that diminish as the reward becomes expected. These data show that the neurotransmitter levels of both, DA and ACh, follow neuronal activity of their respective neurons with a sub-second kinetic. The fast induction of the ACh dip is particularly striking as it suggests fast degradation or diffusion of ACh.

In addition, we observed similar changes in DA and ACh levels to the conditioned stimulus and not the unconditioned stimulus, which occur in parallel over learning. Our data confirm thus that both DA neurons and CINs respond to salient and conditioned stimuli. Moreover, we found that these changes in DA and ACh levels correlate with behavioral responding. However, this correlation was not observed in all tested mice due to the nature of the Pavlovian task. In the Pavlovian task, animals have the possibility to learn an association between the CS+ and reward. Consequently, some animals may show anticipatory responding during the CS+ (head poking the reward port). However, because anticipatory behavior is not fully required to obtain a reward, some of the tested mice did not exhibit anticipatory head poking. Strikingly, the development of a DA and ACh signal over time indicate that these animals are nevertheless learning the stimulus-reward association. In conclusion, these data show that task-evoked changes in ACh and DA levels in mice follow what has been described at the level of neuronal activity level in primates (Joshua et al., 2008; Morris et al., 2004; Schultz et al., 1997)

### D2R inactivation in CINs shortens but does not abolish the cue induced ACh dip

Salient and conditioned stimuli are known to induce pauses in CIN and TAN firing in rodents and primates, however, the dependence on DA and CIN D2Rs for pause induction is widely debated (Aosaki et al., 1994a; Morris et al., 2004; Watanabe & Kimura, 1998; Zhang & Cragg, 2017). Thus, to determine the role D2Rs play in modulating the stimulus induced ACh dip, we pharmacologically blocked D2Rs or selectively ablated Drd2 from CINs and measured ACh and DA levels in the CRF task. We found that D2R blockade or ablation shortened the ACh dip, which we quantified by calculating the dip length. Moreover, in our control mice, D2R blockade also decreased the negative and increased total and rebound AUCs in a dose-dependent manner while the dip amplitude was unaffected. In contrast, D2R blockade had no effect on the stimulus induced ACh dip in ChATDrd2KO mice. This data reveals that the generation of ACh dip is not dependent on CIN D2Rs. Instead, cholinergic D2Rs are important for modulating the length of the stimulus induced ACh dip. Our data provide clarity on the controversial role that DA plays in the regulation of the ACh dip and suggest that the stimulus induced ACh dip *in vivo* is not entirely DA or D2R-dependent as studies in primates and slice physiology studies have suggested (Aosaki et al., 1994a; Ding et al., 2010; Watanabe & Kimura, 1998). Our data further indicates that slice physiology studies in rodents where optogenetic stimulation of DA terminals or caged DA induced CIN pauses are abolished by D2R antagonists or CIN-selective ChATDrd2KO mice are not fully capturing the natural pause (Augustin et al., 2018; Chuhma et al., 2014; Kharkwal et al., 2016; Straub et al., 2014; Wieland et al., 2014).

In addition to the effects on dip lengths, we found that CIN D2Rs also regulate the level of ACh rebound levels, acting as a mechanism to constrain ACh rebounds after the dip. Currently, it is unknown which role the rebound in ACh plays during behavior. Generally, CINs are thought to inhibit spiny projections neurons (SPNs) via nicotinic activation of local interneurons or via muscarinic M_2_/M_4_-mediated inhibition of corticostriatal inputs (English et al., 2012; Faust et al., 2015; Pakhotin & Bracci, 2007; Witten et al., 2010). Thus, a larger dip may lead to disinhibition and higher rebound to a stronger inhibition of SPNs. D2R antagonism decreases the first and enhances the second which may inhibit movement initiation leading to the longer latency in lever pressing. Consistent with this, we observed that the size of the cue induced dip correlates with press latency (the larger the dip the shorter the latency). Surprisingly, this relationship also holds true for lever presses that were performed long after the cue induced ACh signal reverted to normal. This suggests that the cue-evoked dip signals the motivational state of the animal. This finding is consistent with recent inhibition studies in which CIN inhibition in the NAc during Pavlovian to Instrumental Transfer (PIT) enhanced the ability of the pavlovian cue to invigorate behavior (Collins et al., 2019). Note, however, that mice with selective Drd2 ablation do not show a deficit in PIT suggesting a more subtle deficit affecting latencies rather than the level of responding (Gallo et al., 2021).

### DA and ACh correlation during task-dependent behaviors

Our approach to simultaneously image both DA and ACh in the same animal allowed us to examine the relationship between these two neuromodulators within trials. In both C57BL/6J and Drd2^fl/fl^ control mice, we identified a strong negative correlation with the cue-induced DA release leading the ACh dip that is attenuated by D2R antagonism. This strong negative correlation between DA and ACh is significantly reduced in ChATDrd2KO mice compared to Drd2^fl/fl^ controls and ChATDrd2KO mice are unaffected by D2R blockade. We also found a weaker positive correlation with DA leading ACh that is enhanced by D2R antagonism. We believe that this positive correlation reveals the rebound in ACh activity following the dip that is blunted by D2R activation at baseline. This data suggests, as discussed above, that CIN D2Rs not only modulate the ACh dip but also the rebound activity.

### Implications for behavior

What does an altered ACh signal mean for behavior? CIN selective D2R knock out mice learn the Pavlovian task presented in Figure 1 as well as control littermates (data not shown). This suggests that even with a shortened dip mice still can learn cue-reward associations. Similarly, we recently described that enhancing the dip lengths by selective overexpression of D2Rs in CINs of the NAc (D2R-OE_NAcChAT_ mice) did not affect Pavlovian learning but was associated with a deficit in Go/No-Go learning (Gallo et al., 2021). Opposite to what we found in this present study using ChATDrd2KO mice, rebound ACh levels are lower in D2R-OE_NAcChAT_ mice. Thus, relatively higher rebound levels in wild-type control mice may suppress SPN activity and responding. However, rebound activity was significantly greater in incorrect No-Go (press) relative to correct No-Go (withhold) trials questioning this hypothesis. Rather we observed that the ACh dip in control mice was different between Go and No-Go trials, in contrast to what we observed in D2R-OE mice. This difference supports the hypothesis that D2R-OE mice did not learn the switch from the Go to the No-Go component of the task due to reduced contrast in ACh dip information between Go and No-Go contingencies. This is consistent with prior studies reporting that striatal ACh is not necessary for initial learning but is important for behavioral performance when task contingencies change (Aoki et al., 2015; Bradfield et al., 2013; Brown et al., 2010; Favier et al., 2020; Okada et al., 2014; Okada et al., 2017; Ragozzino et al., 2009).

In this present study, we report that D2R antagonism increases lever press latency during CRF in a dose-dependent manner, but this is abolished in ChATDrd2KO mice. Moreover, lever press latency correlated with the AUC of the ACH signal, which was mostly driven by dip size (the larger the dip the shorter the latency) and was observed in trials with latencies longer that the ACh signal lengths. As discussed, this suggests that the cue-evoked dip signals the motivational state of the animal confirming a role of ACh in motivated behavior (Aosaki T, 1994b; Apicella, 2007; Apicella et al., 1991; Collins et al., 2019; Joshua et al., 2008; Kimura et al., 1984; Morris et al., 2004; Nougaret & Ravel, 2015; Ravel et al., 2003; Shimo & Hikosaka, 2001).

In conclusion, our data demonstrate that the stimulus induced ACh dip is multiphasic, encompassing a DA component and a non-DA component. Striatal DA is responsible for confining the temporal boundaries of the ACh dip and preventing rebound excitation via CIN D2R. Notably, we also find a positive correlation between the size of the stimulus induced ACh dip and behavioral responding, which implicates a role for ACh in motivated behaviors. Thus, further dissection of this system will provide a better understanding of the different components of the ACh dip and what each component represents for specific striatal functions and behavior.

## Supplemental figures

**Supplementary Figure 1.**
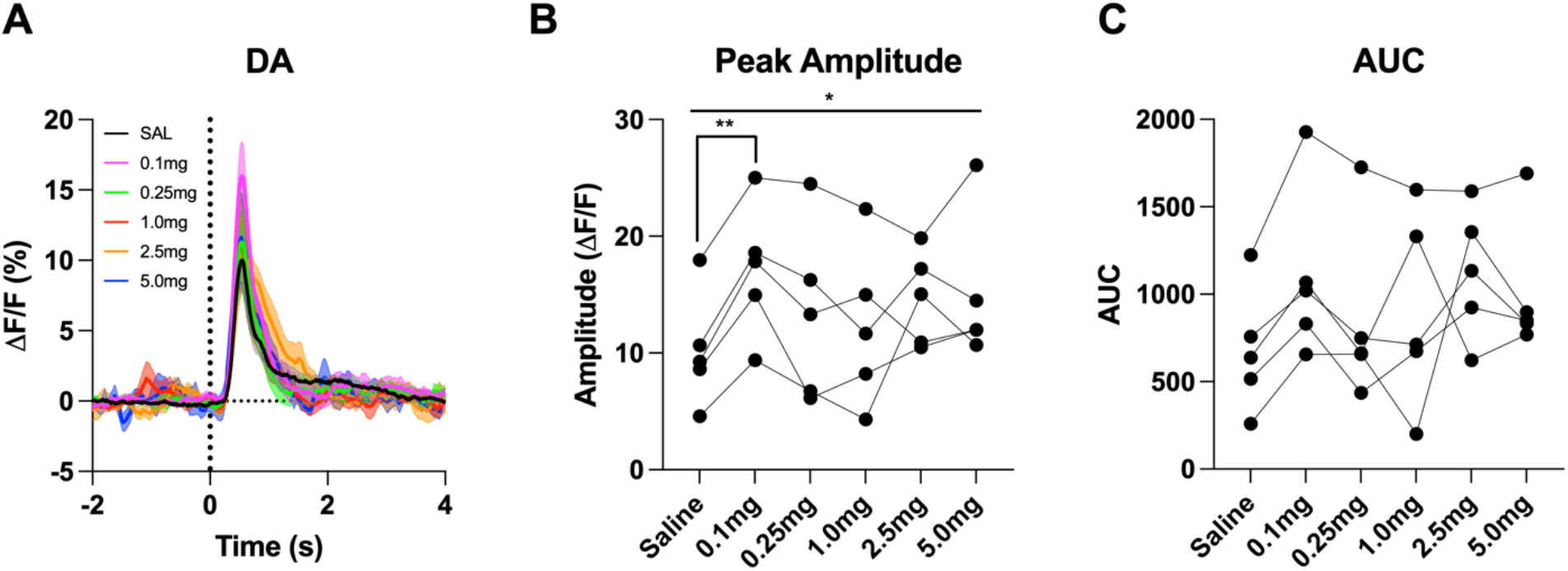
D2R antagonism increases phasic DA release. **(A)** Changes in DA fluorescence (ΔF/F (%)) aligned to lever extension with saline (black) and increasing doses of eticlopride: 0.1 mg/kg (pink), 0.25 mg/kg (green), 1.0 mg/kg (red), 2.5 mg/kg (orange) and 5.0 mg/kg (blue) in C57BL/6J mice. **(B)** Peak amplitude is increased by eticlopride in a dose-dependent manner (F_(2.201, 8.805)_ = 4.268, p = 0.0480). The most prominent increase in peak amplitude is between saline and 0.1 mg/kg (p = 0.0069). **(C)** No overall effect of eticlopride on the AUC (F_(1.440, 5.759)_ = 2.347, p = 0.1807).

**Supplementary Figure 2.**
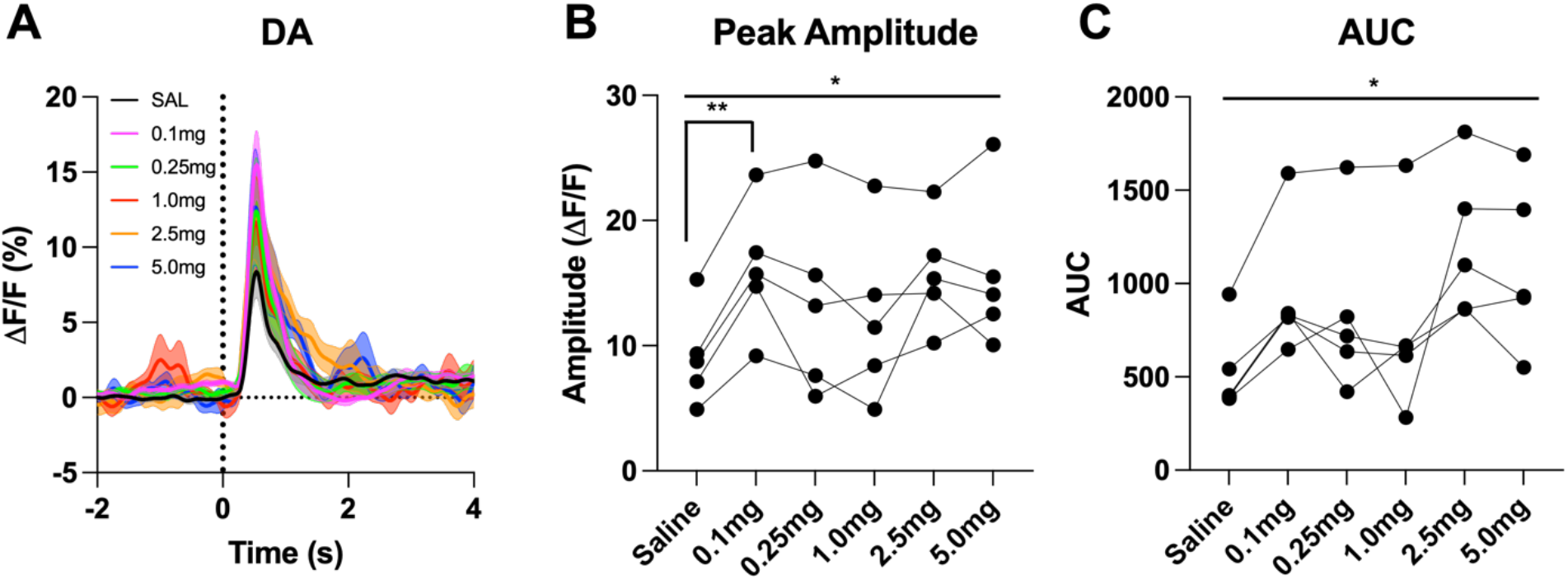
D2R antagonism enhances cue evoked DA release for trials with press latencies > 2s. **(A)** Changes in DA fluorescence (ΔF/F (%)) aligned to lever extension for only trials with press latencies > 2 s with increasing doses of eticlopride in C57BL/6J mice. **(B)** Peak amplitude of DA is increased by eticlopride in a dosedependent manner (F_(2.785, 11.14)_ = 5.804, p = 0.0133) with the most prominent increase between saline and 0.1 mg/kg (p = 0.0099). **(C)** DA AUC is increased by eticlopride in a dose-dependent manner (F_(1.822, 7.288)_ = 6.872, p = 0.0244).

**Supplementary Figure 3.**
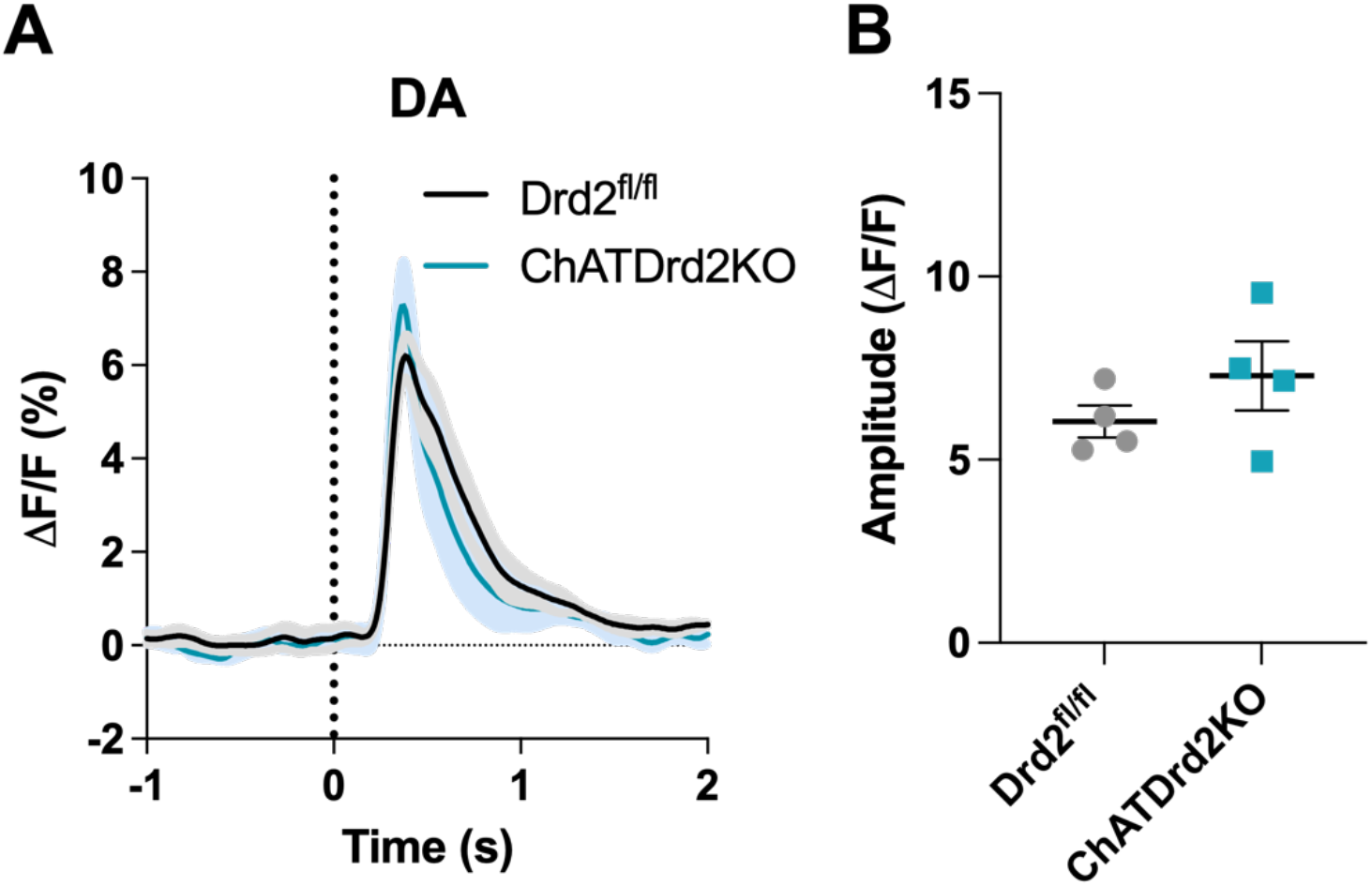
Selective D2R ablation from CINs does not alter cue-evoked DA release for trials with press latencies > 2 s. **(A)** Changes in DA fluorescence (ΔF/F (%)) aligned to lever extension for only trials with press latencies > 2 s for Drd2^fl/fl^ control (black) and ChATDrd2KO (blue) mice, N=4/ genotype. **(B)** The DA peak amplitude is comparable between ChATDrd2KO and Drd2^fl/fl^ control mice (p= 0.2744).

**Supplementary Figure 4.**
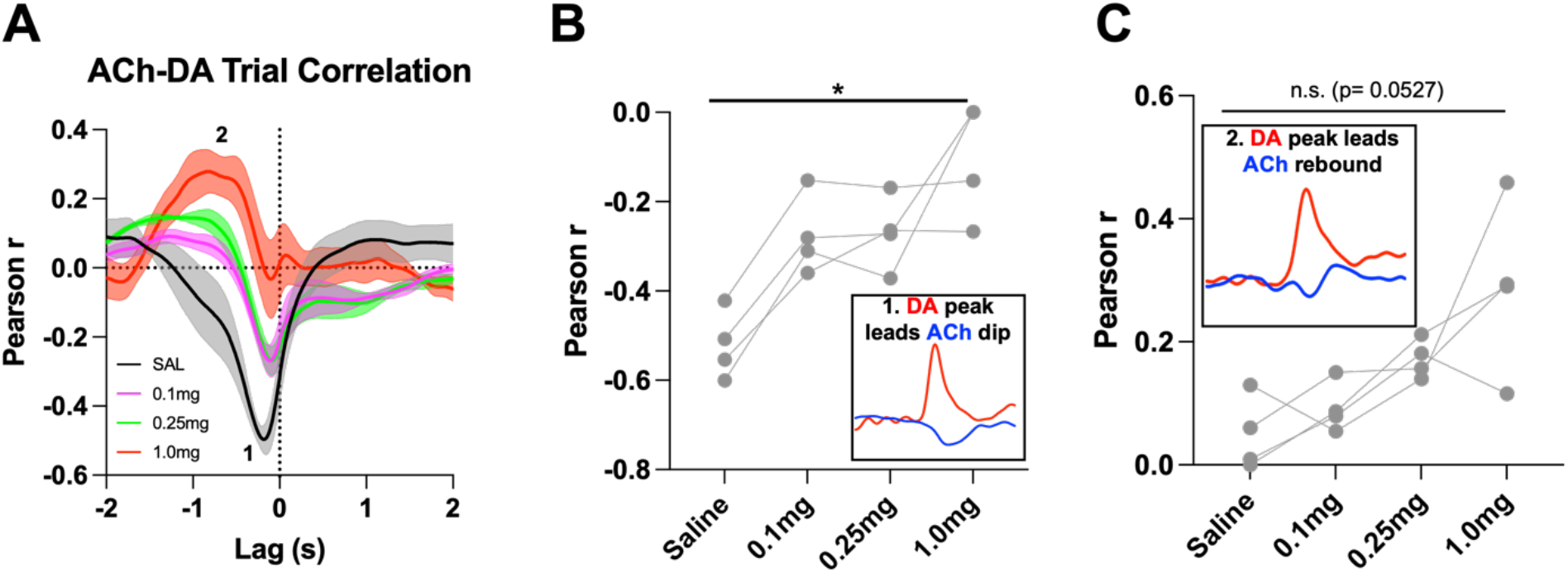
D2R antagonism alters cue-evoked ACh-DA interactions at the lever extension. **(A)** Correlation between ACh and DA during CRF trials with increasing doses of eticlopride in Drd2^fl/fl^ control mice. The ACh signal moved in front of or behind the DA signal to identify points of highest correlation. The first correlation is a negative correlation (1) with ACh lagging DA (Lag= −167.12ms ±18.82 ms) and the second correlation is a positive correlation (2) with ACh lagging DA (Lag= −1.51 s ± 0.138 s) **(B)** The negative correlation with the DA peak leading the ACh dip (inset) is decreased by eticlopride in a dose-dependent manner (F_(1.141, 3.424)_ = 15.48, p = 0.0221). **(C)** The positive correlation with the DA peak leading the ACh rebound (inset) is increased by eticlopride in a dose-dependent manner (F_(1.338, 4.014)_ = 7.088, p = 0.0527).

**Supplementary Figure 5.**
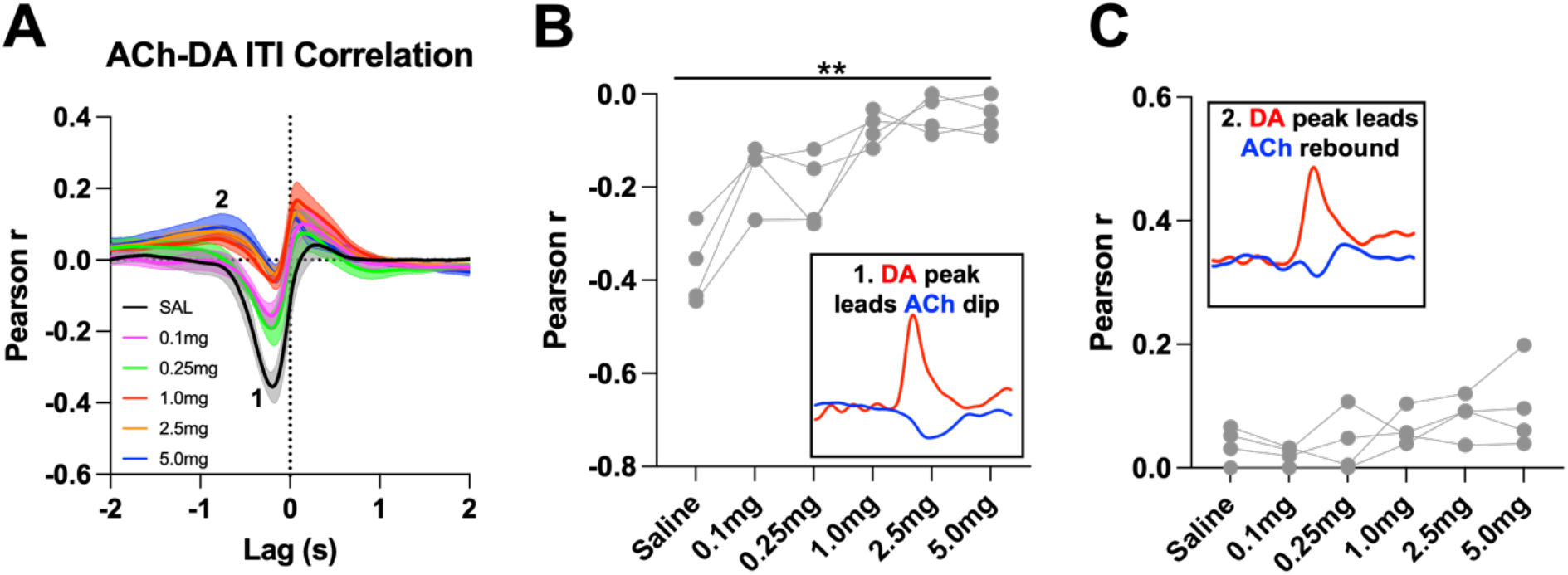
D2R antagonism alters general ACh-DA interactions during the ITI in Drd2^fl/fl^ control mice. **(A)** Correlation between ACh and DA during the ITI with increasing doses of eticlopride in Drd2^fl/fl^ control mice. The ACh signal moved in front of or behind the DA signal to identify points of highest correlation. The first correlation is a negative correlation (1) with ACh lagging DA (Lag= −213.81 ms ± 34.14 ms) and the second correlation is a positive correlation (2) with ACh lagging DA (Lag= −1.39 s ± 0.208 s) **(B)** The negative correlation with the DA peak leading the ACh dip (inset) is dose-dependently reduced by eticlopride (F_(2.102, 6.307)_ = 18.68, p = 0.0021). **(C)** The positive correlation with the DA peak leading the ACh rebound (inset) is not significantly affected by eticlopride (F_(1.276, 3.827)_ = 2.504, p = 0.1966).

**Supplementary Figure 6.**
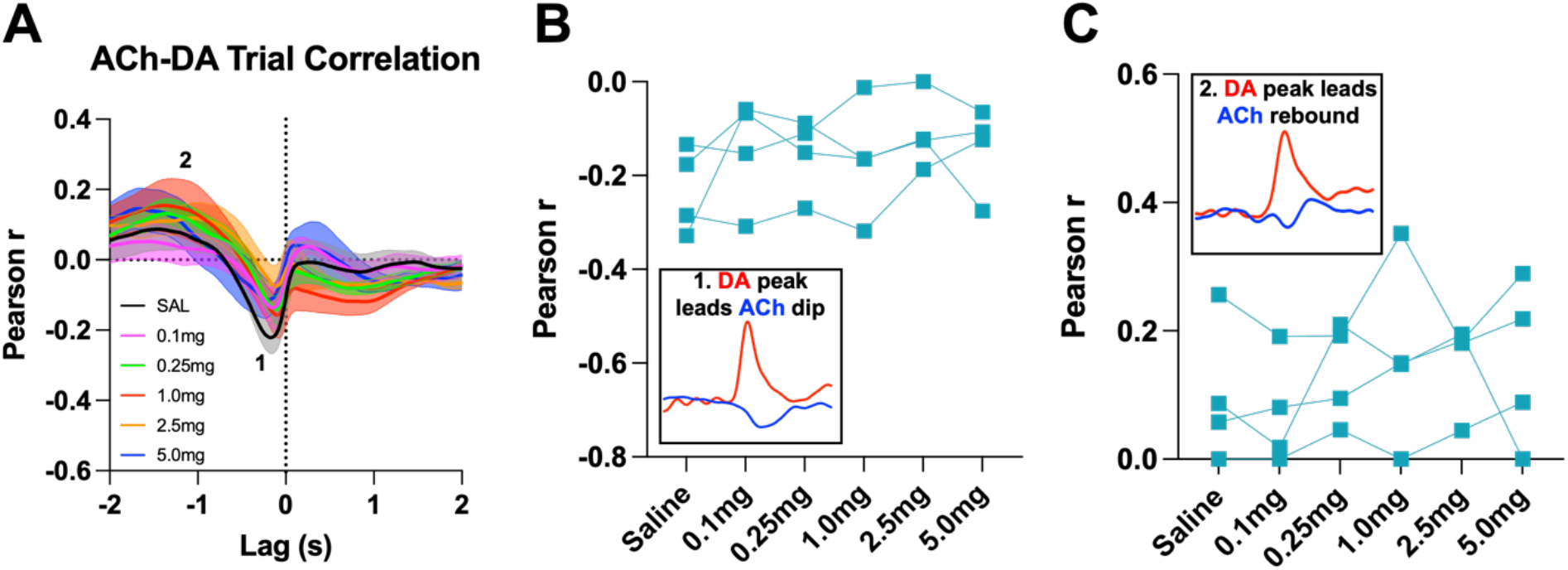
D2R antagonism does not alter cue evoked ACh-DA interactions in ChATDrd2KO mice at the lever extension. **(A)** Correlation between ACh and DA during CRF trials with increasing doses of eticlopride in ChATDrd2KO mice. The ACh signal moved in front of or behind the DA signal to identify points of highest correlation. The first correlation is a negative correlation (1) with ACh lagging DA (Lag= −179.41 ms ± 30.66 ms) and the second correlation is a positive correlation (2) with ACh lagging DA (Lag= −1.635 s ± 0.174 s). **(B)** The negative correlation with the DA peak leading the ACh dip (inset) is not affected by eticlopride (F_(1.720, 5.161)_ = 1.170, p = 0.3682). **(C)** The positive correlation with the DA peak leading the ACh rebound (inset) is not affected by eticlopride (F_(2.016, 6.048)_ = 0.9160, p = 0.4500).

**Supplementary Figure 7.**
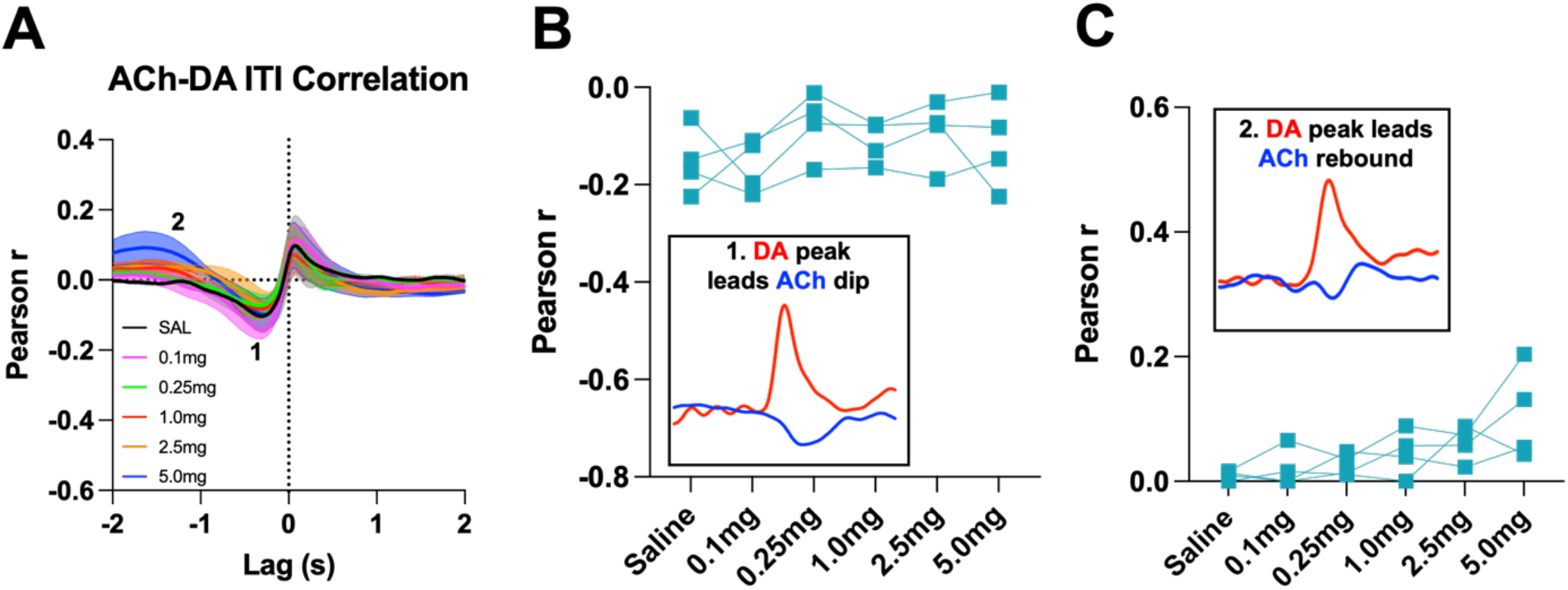
D2R antagonism does not alter general ACh-DA interactions in ChATDrd2KO mice during the ITI. **(A)** Correlation between ACh and DA during CRF trials with increasing doses of eticlopride in ChATDrd2KO mice. The ACh signal moved in front of or behind the DA signal to identify points of highest correlation. The first correlation is a negative correlation (1) with ACh lagging DA (Lag= −282.62 ms ± 93.81 ms) and the second correlation is a positive correlation (2) with ACh lagging DA (Lag= −1.33 s ± 0.189 s) **(B)** The negative correlation with the DA peak leading the ACh dip (inset) is not affected by eticlopride (F_(1.508, 4.523)_ = 1.366, p = 0.3275). **(C)** The positive correlation with the DA peak leading the ACh rebound (inset) is not affected by eticlopride (F_(1.747, 5.240)_ = 4.839, p = 0.0669).

